# Entropy predicts sensitivity of pseudo-random seeds

**DOI:** 10.1101/2022.10.13.512198

**Authors:** Benjamin Dominik Maier, Kristoffer Sahlin

## Abstract

In sequence similarity search applications such as read mapping, it is desired that seeds match between a read and reference in regions with mutations or read errors (seed sensitivity). *K*-mers are likely the most well-known and used seed construct in bioinformatics, and many studies on, *e*.*g*., spaced *k*-mers aim to improve sensitivity over *k*-mers. Spaced *k*-mers are highly sensitive when substitutions largely dominate the mutation rate but quickly deteriorate when indels are present. Recently, we developed a pseudo-random seeding construct, strobemers, which were empirically demonstrated to have high sensitivity also at high indel rates. However, the study lacked a deeper understanding of why. In this study, we demonstrate that a seed’s entropy (randomness) is a good predictor for seed sensitivity. We propose a model to estimate the entropy of a seed and find that seeds with high entropy, according to our model, in most cases have high match sensitivity. We also present three new strobemer seed constructs, mixedstrobes, altstrobes, and multistrobes. We use both simulated and biological data to demonstrate that our new seed constructs improve sequence-matching sensitivity to other strobemers. We implement strobemers into minimap2 and observe slightly faster alignment time and higher accuracy than using *k*-mers at various error rates.

Our discovered seed randomness-sensitivity relationship explains why some seeds perform better than others, and the relationship provides a framework for designing even more sensitive seeds. In addition, we show that the three new seed constructs are practically useful. Finally, in cases where our entropy model does not predict the observed sensitivity well, we explain why and how to improve the model in future work.

## 1 Introduction

Short *k*-length substrings of a sequence, often referred to as *k*-mers, are widely used for sequence comparison in bioinformatic applications. A *k*-mer that is shared by two sequences implies an identical region of size *k*, and with appropriate length on *k*, we may detect similar but non-identical regions through shared *k*-mers. Some of the reasons that *k*-mers are often used for sequence similarity detection is because they are fast to construct, and due to their fixed length easy to represent, store, and query, *e*.*g*., with hash tables, or more succinct data structures such as bloom filters [5], the FM-index [15], and many more [32]. As *k*-mers indicate shared sequences, they are often used as markers, or *seeds*, indicating regions for more extensive similarity comparison, *e*.*g*., through pairwise alignment.

With the broad use of *k*-mers as seeds, several limitations have also been identified. For example, *k*-mers are sensitive to mutations. If *k* is too small, we may obtain many redundant hits, *e*.*g*., due to repeats). On the other hand, a too large *k* may destroy all matches (low *sensitivity*) in mutation-dense regions or error-prone reads. Detailed modeling of *k*-mers’ sensitivity to substitutions at different rates was performed in [3]. Some studies have proposed altering the underlying biological sequence to reduce the mutation rates with, e.g., homopolymer compression [2] or modifying the mutation distribution using more advanced sequence transformations [4]. However, most work has been aimed at increasing seed sensitivity and lower seed repetitiveness of *k*-mers by proposing alternative *seed constructs*.

### 1.1 Other seed constructs

Some approaches aim to alleviate repetitiveness issues in downstream analysis by dynamically extending the k-mers to provide a less redundant set of matching seeds such as Maximal Exact Matches (MEMs), Maximal Unique Matches (MUMs) [11], MCAS [25], and Context-aware seeds [50]. These seeding constructs have been referred to as dynamic seeds [40] as they are neither fixed in length nor in the number of CPU cycles for their construction. There are also seeding constructs known as *subsampling methods* that aim to use only a subsample of *k*-mers as seeds due to their redundant nature using, *e*.*g*., minimizers [36] or later subsampling techniques [10,19,13,51,12,20]. For an extensive study of subsampling techniques, see [42].

To overcome the issue of requiring only exact matches, spaced seeds (or spaced *k*-mers) [30], covering template families [21], indel seeds [31], and SimHash-based constructs [8] such as permutation-based seeds [28] or BLEND [18] have been proposed that, particularly, tolerate substitutions. Covering template families and indel-seeds are also designed to match over small indels, and are based on extracting several fixed-pattern seeds per query position. For example, in [21], a combination of patterns is chosen to provide a guarantee that at least one seed matches. The required number of extracted seeds increases with indel size. Seeds that do not require an identical sequence to match are often called *fuzzy seeds*. In applications where substitutions are frequent, spaced *k*-mers have had practical success and are used in several state-of-the-art applications such as in the general sequence similarity search software BLAST [1], and for metagenomic classification [7] and long-read mapping [43].

#### Strobemers

Recently, we introduced a new class of fuzzy seed constructs, *strobemers* [37], tolerant to substitutions and indels. Strobemers expand on the ideas of neighbouring minimizer pairs [9,41] and *k*-minmers [14]. Strobemers are constructed by linking together a set of smaller *k*-mers and can be constructed with several different methods to link the *k*-mers (minstrobes, randstrobes, hybridstrobes), yielding different properties. Due to the pseudo-random sampling process of strobemers, we will in this work refer to strobemer seeds as *pseudo-random* seeds, not to be confused with other types of fuzzy seeds that can have a non-random sampling process, *e*.*g*., SimHash based on Locality Sensitive Hashing techniques [8]. It was shown that strobemers could offer higher sensitivity and lower repetitiveness over *k*-mers, and they have been used for short-read mapping [39], long-read overlap detection [18], and transcriptomic long-read normalization [33].

### 1.2 Previous work on seed sensitivity

In spaced seed literature, seed sensitivity has been extensively studied. Typically, when using seeds, an alignment is triggered if a certain number of seeds match in a region, *e*.*g*., through requiring either multiple *hits* [6] (seed matches) or a single hit [26] in a region. One drawback of requiring multiple hits is that a threshold does not distinguish highly overlapping hits from disjoint ones. For this reason, seed coverage (union of matching positions in a region) has been proposed [34].

The main conclusion in spaced seed literature is that many highly overlapping seeds are redundant, uninformative, and can lead to unnecessary computations for sequence matching applications. Typically, the aim is to select a set of seed patterns consisting of fixed and wildcard positions that overlap or correlate as little as possible. Related work on minimizing the overlap of hits has been studied in the form of clump statistics [44] or *overlap complexity* [24]. In addition, there are other theoretical studies of seed sensitivity quantifying the correlation between seeds [27] or using generating functions from analytical combinatorics [16], which have also been used in practice to select suitably spaced seeds when mapping short reads [17].

### 1.3 Our aim

The aforementioned spaced-seed studies are all based on seeds with a fixed sampling pattern (*k*-mers and spaced *k*-mers). That is, the sampling decision has no pseudo-random behaviour after the seed pattern has been chosen. Flexible gapped seeds such as strobemers employ a pseudo-random sampling decision based on the underlying sequence. These seeds are a new class of fuzzy seeds that are more tolerant to indels and open up a venue to study how randomness influence seed overlap, correlation, and sensitivity. Also, while there are spaced seed studies focusing on optimal seed selection of a single seed pattern [22], most aforementioned spaced seed studies are centered around selecting a set of spaced seeds with complementary properties. We focus on seed efficacy restricted to one seed per position as is used for *k*-mers, for query efficacy purposes.

Inspired by spaced *k*-mer literature that has focused on understanding the mechanics of high sensitivity seeds, we aimed to find why pseudo-random seeds such as strobemers achieve beneficial properties such as high sensitivity and low repetitiveness. We focused on seed constructs that, similarly to *k*-mers, (1) produce one seed per position, (2) always sample *k* fixed positions in a window of *w* positions (i.e., no dynamic size construction such as MEMs or MCAS), and (3) require only a single look-up in the index for match detection. Here, we call these seeds (*k, w*)-seeds. For example, *k*-mers, spaced *k*-mers, and strobemers are all valid (*k, w*)-seeds. Constraint 1 allows for fair sensitivity and repetitiveness benchmarking. Constraint 2 ensures that seed construction is constant across string composition for easy benchmarking (i.e., not sensitive to repeats as dynamic seeds). Constraint 3 ensures that, after construction, the seeds are equally fast for sequence similarity comparison (e.g., through a hash table look-up).

### 1.4 Paper outline and our contributions

In section 2.1, we state the notation and preliminaries. In section 2.2, we describe pseudo-random seeds and formalize the notion of (*k, w*)-seeds. In section 2.3, we formalize the objective of a good seed for sequence similarity search, namely a seed that has high sensitivity and low repetitiveness. In sections 2.4 and 2.5, we discuss the main finding in this paper, namely that the entropy (randomness) of a seed can be used as a predictor of seed sensitivity. Specifically, in most cases, higher entropy seeds have a higher probability of generating at least one match in a region that has undergone mutations. We provide a model to calculate the entropy of a seed. In section 2.6 we present three new seed constructs that we call *mixedstrobes, altstrobes*, and *multistrobes*.

In section 3.1, we empirically verify that, in most cases, a higher seed entropy results in higher sensitivity for various seed constructs. We also discuss the limitations of our entropy model. In sections 3.2 we evaluate how our proposed new strobemer seeding constructs fares to randstrobes and *k*-mers in an actual sequence matching scenario with the four different metrics used in [37]. Of immediate practical importance, we find that the three constructs can all improve over the currently best-known strobemer construct (randstrobes). Section 3.3 further explore the performance of seeds (including spaced *k*-mers) at different substitution rates. Section 3.4 discuss time and memory requirements for constructing the seeds in C++, and section 3.5 discuss the results of implementing strobemer seeds in minimap2. Even though minimap2 performs subsampling of seeds which distorts the entropy and, thus, our sensitivity predictions, we observe faster alignment time (up to 30%) and slightly higher sensitivity (0.2%) than using *k*-mers of the same size as seeds. Section 4 and 5 discuss the results and describes future direction on how to use our entropy for seed design. In summary, our work presents three new state-of-the-art seeding constructs and our insight and model can be used to design even better seeds than those proposed in this study.

## 2 Methods

### 2.1 Definitions and preliminaries

#### Notation

We define a *subsequence* of a string as a set of ordered nucleotides that can be derived from the string by removing some or no elements while keeping the order of the remaining elements. If all letters in the subsequence are consecutive, we refer to it as a substring. We use 0-indexed notation for indexing sequences or strings, and we write *S*[*i* : *j*], *i < j*, to refer to a substring of a string *S* starting at position *i* and ending at *j* but not including the character at *j*. That is, the start index is inclusive, but the end index is exclusive. We let the | · | operator denote the length of strings, *e*.*g*., |*S*[*i* : *j*]|= *j*− *i*. We let the + operator denote string concatenation if applied to strings. Finally, we use *h* to denote a hash function mapping strings to integers.

#### Strobemers

Strobemers are seeds that consist of *n >* 1 *ℓ*-mers (strobes)[37]. The first strobe *s*_1_ is the *ℓ*-mer at the position where the seed should be extracted. The subsequent strobes *s*_2_, … *s*_*n*_ are chosen from downstream windows defined by a lower (*w*_*min*_) and upper (*w*_*max*_) offset to its respective previous strobe’s window. Hence, strobemers are characterized by the number of strobes (*n*), the length of the strobes (*ℓ*) and the window constraints (*w*_*min*_ and *w*_*max*_). Three methods to construct downstream strobes *s*_2_, …, *s*_*n*_ was given in [37], ; minstrobes, hybridstrobes and randstrobes. For minstrobes, strobe *s*_2_ is simply the minimizer [36] in the window [*w*_*min*_, *w*_*max*_] downstream from the first strobe. Hybridstrobes partitions [*w*_*min*_, *w*_*max*_] into *x* sub-windows, and picks as *s*_2_ the minimizer in sub-window *h*(*s*_1_)%*x* (ordered 0 to *x* − 1). Randstrobes, the most effective seed in [37] selects as *s*_2_ the *ℓ*-mer *s*^*′*^ in the window [*w*_*min*_, *w*_*max*_] that minimizes a hash function *h*(*s*_1_ + *s*^*′*^) (although there are variations to string concatenation, see [37]).Further downstream strobes if *n >* 2 are sampled analogously to *s*_2_. This study will mostly consider strobemers with *n* = 2.

#### E-hits of seeds

We will in our assessment of seeds need a notion of seed repetitiveness, and we will use E-hits for this. The definition of E-hits was given in [39] and is a measure of how repetitive the seeds in a query sequence are, on average, in a reference dataset. More specifically, E-hits computes the expected number of hits that a seed matched to the reference will have, given that the seed is error free and come from a query uniformly sampled from the reference. E-hits can be calculated for any seeding mechanism and reference dataset. For a given reference dataset, let *N* be the total number of seeds sampled, *M* the total number of distinct seeds sampled, and *z*_*i*_ be the total number of times the distinct seed *i* (1 ≤*i* ≤*M*) is sampled. Then E-hits is computed as follows

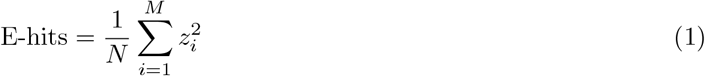

### 2.2 Pseudo-random seeds

At a high level, a *fuzzy seed* is a subsequence extracted from a string *S* that is guaranteed to match another fuzzy seed extracted from another string *T* if *S* = *T* and may match *T* if *T* is similar but not identical to *S*. In this work, we consider *pseudo-random seeds*. With a pseudo-random seed, we mean a fuzzy seed where there is also pseudo-randomness in the construction process. Pseudo-random seeds can be considered a subclass of fuzzy seeds because there exist fuzzy seed constructs where the construction process has no randomness, *e*.*g*., spaced *k*-mers or SimHash [8]. Furthermore, a pseudo-random seed will be fuzzy implied by the multiple choices in the construction process. Hybridstrobes and randstrobes are examples of pseudo-random seeds.

For convenience, we use the following general notation for the construction of seeds. Let *k* and *w* be two positive integers with *k* ≤ *w* where *k* denotes the number of distinct positions sampled in a substring of length *w* in a string *S*. Let *f* (*i, k, w, S*, *) denote some function that starts at position *i* in *S* and extracts a subsequence of characters at *k* distinct positions in the substring *S*[*i* : *i* + *w*] using only the information in *S*[*i* : *i* + *w*]. We use the final argument *∗* to denote any seeding specific parameters which may include, e.g., the sampling pattern for spaced *k*-mers, or parameters (*n, l, w*_*min*_, *w*_*max*_) for strobemers. For the seed to be a pseudo-random seed, *f* needs to employ a pseudo-random sampling decision. Other seeds such as *k*-mers and spaced *k*-mers can be produced under this description of *f* (when *f* is a fixed sampling pattern) but they are not pseudo-random seeds. We will refer to seeds constructed by any *f* with parameters *k* and *w* as a (*k, w*)-seed.

#### Constraints on *f*

We impose the following three basic constraints on *f* to be viable for sequence matching.

C1 *f* produce the same (*k, w*)-seeds for two strings *S* and *T* if *S* = *T*.

C2 *f* produce valid (*k, w*)-seeds ∀*S, S* ∈ Σ^*∗*^.

C3 at most one seed is produced per position in a sequence.

C1 and C2 are necessary for sequence matching. An example of a construct that violates C2 is “sample the position if the letter is A or C” because there may not be enough A’s and C’s in the window. C3 limits querying to at most one lookup per position making the constructs efficient. We have intentionally described *f* in a general fashion in order to encompass more general construction techniques. For example, a *k*-mer would deterministically sample the first *k* nucleotides (nt), regardless of the size of *w*. A randstrobe [37] with parameters (2,15,25,50) is a valid (30, 64)-seed because the maximum span of sampled positions in the randstrobe is *w* = 50 + 15 − 1 = 64 and *f* would sample 15nt at *S*[*i* : *i* + 15] as the first strobe, then sample the next strobe starting somewhere in *S*[*i* + 25 : *i* + 50].

### 2.3 Objectives for sequence similarity detection

Our objective is similar to what was sought but not thoroughly formulated in [37]. We state the objectives in precise terms here. Let two strings *S* and *T*, each of length 2*w* have an edit distance *m* ≥ 0 to each other. Let *N*_*m*_(*k, w*) be the number of seed matches from the first *w* consecutive (*k, w*)-seeds constructed from *S* and *T* (see Fig. 1A). We desire a function *f* that extract (*k, w*)-seeds such that:

**Fig. 1:**
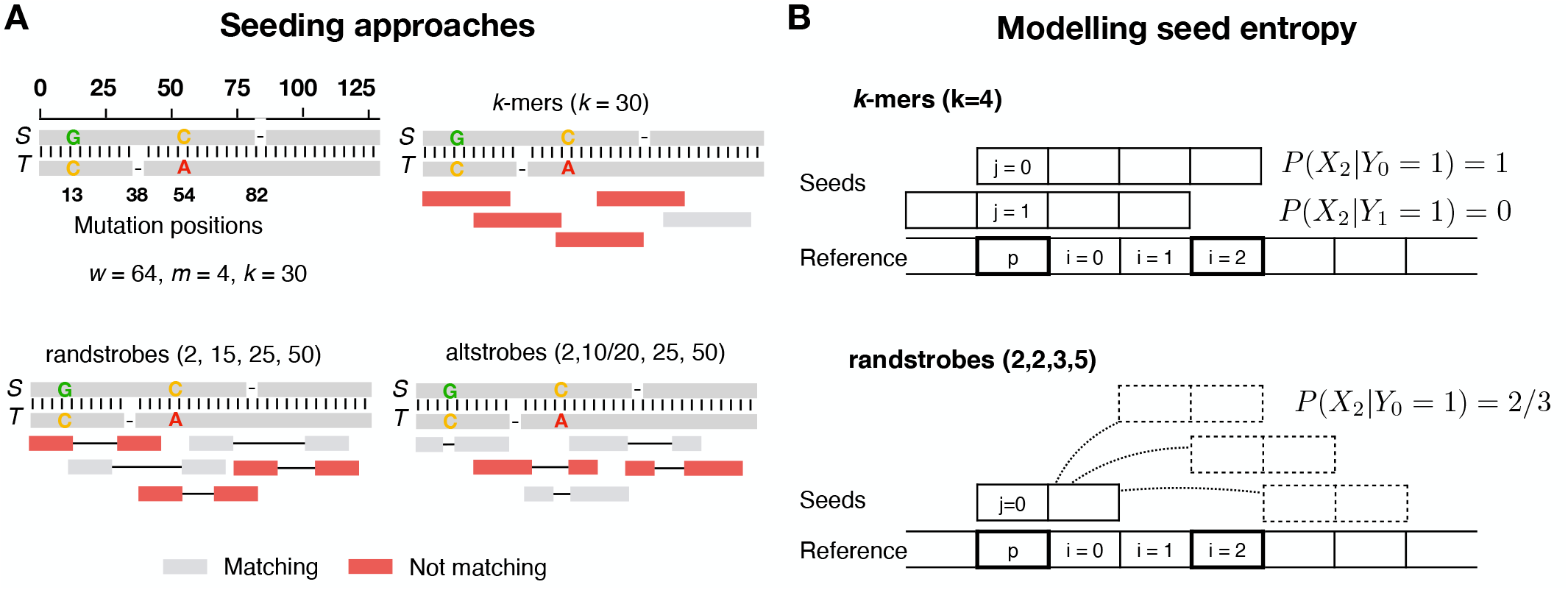
Sampling of k-mers, randstrobes and altstrobes. Panel A shows a scenario where *k*-mers, randstrobes and altstrobes are sampled over two similar sequences *S* and *T* different by four mutations. Only a subset of five seeds in the region is shown for clarity. In the example, the seed constructs all sample 30 fixed positions in a window of *w* = 64 nucleotides which is the maximal span of a randstrobe with parameters (2,15,25,50). The mutations are distributed such that they destroy all shared k-mers in the region, and most of the randstrobes. Altstrobes have the possibility to sample k-mers of two different lengths at each site, which allow them a higher probability to match between mutations. Panel B illustrates our modelling of seed entropy for seed construct *k*-mers and randstrobes. In the case of *k*-mers, there is no pseudo-randomness and therefore all probabilities are either 0 or 1, leading to an entropy of 0. Under uniform hashing, the randstrobes will have a probability of 2/3 of covering position *i* due to the three possible sampling positions of the second strobe. Boldfaced squares indicate the positions considered for the computation of probabilities shown in the figure.

O1 *P* (*N*_*m*_(*k, w*) *>* 0) is as large as possible ∀*m* ≥ 0.

O2 The E-hits metric [39] for *f* is as small as possible.

O1 relates to seed sensitivity, and O2 relates to seed repetitiveness. The formulation of *N*_*m*_(*k, w*), namely to only consider the first *w* seeds in a region of 2*w* for short strings, may seem unfair to *k*-mers. This is because *K*-mers can produce additional hits between *S* and *T* from the last *w*−*k* seeds at the ends of *S* and *T* (see Fig. 1A). This advantage is present at the end of each sequence, or at the end of every alignment site in case of split alignments by large indels or rearrangements. We aim to model a scenario where sequences are substantially longer than the extra *w*−*k* seeds in the ends, *e*.*g*., as for long reads. Therefore, O1 reflects all regions but the *w*-long end region of sequences.

### 2.4 Randomness influence seed sensitivity

In [37], it was shown that, e.g., randstrobes and hybridstrobes had higher sensitivity (better at finding matches) than *k*-mers, spaced *k*-mers and minstrobes (Fig. 1 and Tab. 1 in [37]). These two constructs have, unlike the rest, a pseudo-random component in how they select the next strobe, creating a seemingly more random seed coverage distribution (see Fig. 1A). In turn, randstrobes had a higher sensitivity than hybridstrobes. By definition of the constructs, the randomness in sampled positions for the second strobe is higher for randstrobes than for hybridstrobes. Therefore, it stands to reason that something in the randomness of the sampled positions of a seed may be positively correlated with seed sensitivity. To formulate it in a verifiable question, does the probability of having at least one match in a region, *P* (*N*_*m*_(*k, w*) *>* 0), increase with higher seed sampling entropy? For the impatient reader, the answer to this question will be, no, not in all cases, but it is in general a good predictor.

### 2.5 Modelling entropy of a pseudo-random seed

We estimate the entropy of a seed as follows. Consider a position *p* on a string *S* with *w* − 1 ≤ *p* ≤ |*S*|− *w, i*.*e*., *p* is not close to the boundaries. For any given seed, let *Y*_*j*_ ∈ 0, 1 be the binary variable denoting if position *j* ∈ [0, *k* − 1] on the seed covers *p* on *S* (*Y*_*j*_ = 1). Let *i* be an indexing integer on the reference starting at *i* = 0 for the position immediately downstream to *p*, and *X*_*i*_ be the binary variable describing the event that position *i* is sampled by the same seed (Fig.1B). Then *X*_*i*_|*Y*_*j*_ describes the event that position *i* was sampled given that *p* was covered by position *j* on the seed, and *P* (*X*_*i*_|*Y*_*j*_) denotes the probability of this event. The conditional entropy of a seed sampling any downstream position given that *p* was sampled by the seed, *X*|*Y*, is computed as

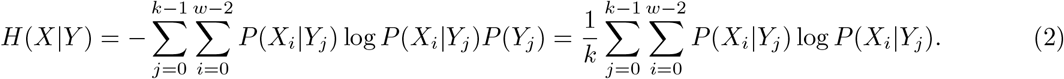

Here *P* (*Y*_*j*_) = 1*/k* by the assumption that any position *j* on the seed is equally likely to cover the reference position *p* if we pick a position *p* at random. Also, variable *i* only needs to be summed up to *w* −2 as positions further downstream will have a probability of 0. The probability, *P* (*X*_*i*_|*Y*_*j*_) is specific to the seed construct but can, for strobemers, be structured up into cases and is relatively straightforward to compute. We provide example computations for *k*-mers, randstrobes, altstrobes, mixedstrobes, and multistrobes in Suppl. section S2. All seed constructs without pseudo-randomness such as k-mers and spaced k-mers have an entropy of 0 according to equation 2, as *P* (*X*_*i*_|*Y*_*j*_) is either 0 or 1.

Our entropy measure cannot estimate entropy for seed constructs that pass information (are correlated) between neighboring seeds for pseudo-random decisions. Example of such a seed construct is hybridstrobes which use minimizers [36], which can be shared between neighboring windows. Nevertheless, we will see that the estimate will be a useful predictor for randstrobes and other pseudo-random seed constructs that we introduce in this study. Finally, our model is agnostic to the underlying error pattern, e.g., the relative fraction of substitutions to indels. It is known that some patterns such as spaced *k*-mers perform well when substitutions are more frequent than indels.

### 2.6 Alternative strobemer constructs

Based on our intuition that randomness would improve sensitivity and our designed model for estimating entropy, we wanted to explore whether altering various parameters in the original strobemer constructs (proposed in [37]) would lead to higher entropy and, therefore, higher sensitivity. We here propose three alternative seed constructs to the strobemer seed-family, mixedstrobes, altstrobes, and multistrobes. We will later demonstrate that these seed constructs, for some parametrizations, can yield higher sensitivity than randstrobes which was the most sensitive seed proposed in [37]. Notably, the parametrizations that receive higher sensitivity than randstrobes also receive higher entropy in our model, although the reverse is not always true.

#### Mixedstrobes

Mixedstrobes samples either a *k*-mer or a strobemer at a specified fraction. Any strobemer may be sampled, but we will only consider randstrobes here. We parameterize mixedstrobes as (*n, ℓ, w*_*min*_, *w*_*max*_, *q*), where *n* is the number of strobes, *ℓ* is the strobe length, *w*_*min*_ and *w*_*max*_ the minimum and maximum downstream offset to last window, and *q* the strobemer fraction. Whether a strobemer or a *k*-mer is seeded depends on the hash value of the first strobe *h*(*S*[*i* : *i* + *ℓ*]) and the user-defined strobe fraction *q*. The strobe fraction *q* is represented as numerator *N* and a denominator *D* (e.g., *q* = 0.6 is represented as *N* = 60 and *D* = 100) so that

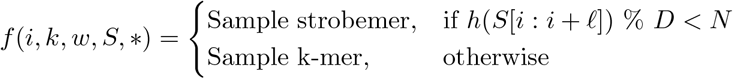

Sample k-mer, otherwise The full pseudocode to construct mixedstrobes is given in Algorithm 1 in Suppl. section S1.

#### Altstrobes

Altstrobes are modified randstrobes where the strobe length alternates (hence altstrobes) between shorter and longer strobes. For example, instead of having two strobes of length *k/*2 as implemented in randstrobes of order 2, altstrobes of order 2 consist of one short strobe *k*_*s*_ and one longer strobe *k*_*l*_, with |*k*_*s*_| + |*k*_*l*_| = *k*. We parameterize altstrobes as (*n*, |*k*_*s*_|, |*k*_*l*_|, *w*_*min*_, *w*_*max*_). We refer to sampled altstrobe with *n* = 2 as (|*k*_*s*_|, |*k*_*l*_|) or (|*k*_*l*_|, |*k*_*s*_|), depending on if the short strobe was used first or second, respectively. We decide the length of the first strobe based on the hash value of the substring of length |*k*_*s*_| (*i*.*e*., the potential first strobe). Specifically,

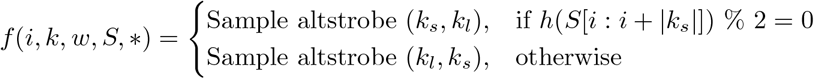

The sampled strobe length is decided by the hash value of the shorter strobe. Otherwise, mutations within the positions [*k*_*s*_, *k*_*l*_] downstream from the start position may lead to seeds being sampled differently between two sequences, which leads to unnecessary seed mismatches. The sampling of the second strobe is performed identically to randstrobes in a downstream window specified by *w*_*min*_ and *w*_*max*_ as described in [37].

For fair benchmarking to other strobemer seeds, we implement two evaluation-specific constraints on altstrobes. Firstly, *n* has to be even to guarantee seeds with the same number of sampled positions. Secondly, to guarantee that all altstrobe seeds are (*k, w*)-seeds, we adjust the sampling window offset depending on if it is the long or short strobe we sample first. Specifically, we let *k*_*l*_ in altstrobe (*k*_*s*_, *k*_*l*_) be sampled from [*w*_*min*_ −(*k*_*l*_ −*k*_*s*_)*/*2, *w*_*max*_ −(*k*_*l*_ −*k*_*s*_)*/*2] and *k*_*s*_ in altstrobe (*k*_*l*_, *k*_*s*_) be sampled from [*w*_*min*_ +(*k*_*l*_ −*k*_*s*_)*/*2, *w*_*max*_ + (*k*_*l*_ − *k*_*s*_)*/*2]. These constraints are only implemented for controlled benchmarking. The full pseudocode to construct altstrobes is given in Algorithm 2 in the Suppl. section S1.

#### Multistrobes

Multistrobes are generalized altstrobes where strobe lengths are selected in a range of lengths. They are parameterized identically to altstrobes as (*n*, |*k*_*s*_|, |*k*_*l*_|, *w*_*min*_, *w*_*max*_) with |*k*_*s*_| + |*k*_*l*_| = *k*. However, unlike altstrobes they can sample any length of a strobe in [|*k*_*s*_|, |*k*_*l*_|]. Similar to, but more generally expressed than altstrobes, the strobe length of the first sampled strobe *k*_1_ in multistrobes is given by |*k*_1_| ≐ |*k*_*s*_|+ *h*(*S*[*i* : *i* + |*k*_*s*_|])%(|*k*_*l*_| − |*k*_*s*_| + 1). The second strobe length is then given the length |*k*_2_| ≐ *k* − |*k*_1_| and is sampled identically to altstrobes in a downstream window. We employ the same evaluation-specific constraints as described for altstrobes, for fair benchmarking to other strobemers.

For small *k*_*s*_, uniform sampling of lengths is not possible. For example, for |*k*_*s*_| = 1, only 4 possible hash values can be produced, which may be smaller than |*k*_*l*_ − *k*_*s*_ + 1|. Similar but not as extreme effects are present for |*k*_*s*_| 2 and 3, especially if several of the few possible hash values map to the same lengths. We employ some implementation tricks to keep a high uniformity despite low *k*_*s*_ given in Suppl. Section S1.3. The full pseudocode to construct multistrobes is given in Algorithm 3 in Supplementary materials.

## 3 Results

### 3.1 Empirically testing the link between entropy and sensitivity

As a first experiment to build intuition, we computed the *P* (*N*_*m*_(*k, w*) *>* 0) for *m* ∈ [1, 25] for *k*-mers, minstrobes, hybridstrobes, randstrobes, mixedstrobes, altstrobes, and multistrobes. We chose *w* = 64 because it corresponds to the total span of the order 2 strobemer seeds parametrized with (2,15,25,50) in [37]. The result is illustrated in Figure 2A. If we sum over *m* as a proxy for sensitivity across mutation rates, we get the estimated summed sensitivity to 9.51, 9.96, 12.28, 12.54, 12.87, 12.95, 13.04 for minstrobes, *k*-mers, hybridstrobes, randstrobes, mixedstrobes (*q*=0.9), altstrobes (*k*_*s*_=10), and multistrobes (*k*_*s*_=5) respectively. Such sensitivity comes at roughly no cost in seed uniqueness (Fig. 2B). We will, in later experiments, see that *k*-mers, randstrobes, mixedstrobes (*q*=0.9), altstrobes (*k*_*s*_=10), and multistrobes (*k*_*s*_ = 5) are also sorted in increasing entropy. As previously stated, our model cannot estimate entropy for minstrobes and hybridstrobes because of shared information. However, intuitively it makes sense for hybridstrobes to have higher entropy than *k*-mers but lower entropy than randstrobes, which is where it places in terms of sensitivity. Also, minstrobes perform worse than other strobemers because it has no sampling pseudo-randomness (similar to *k*-mers). This result builds intuition and indicates that higher entropy implies higher sensitivity.

**Fig. 2:**
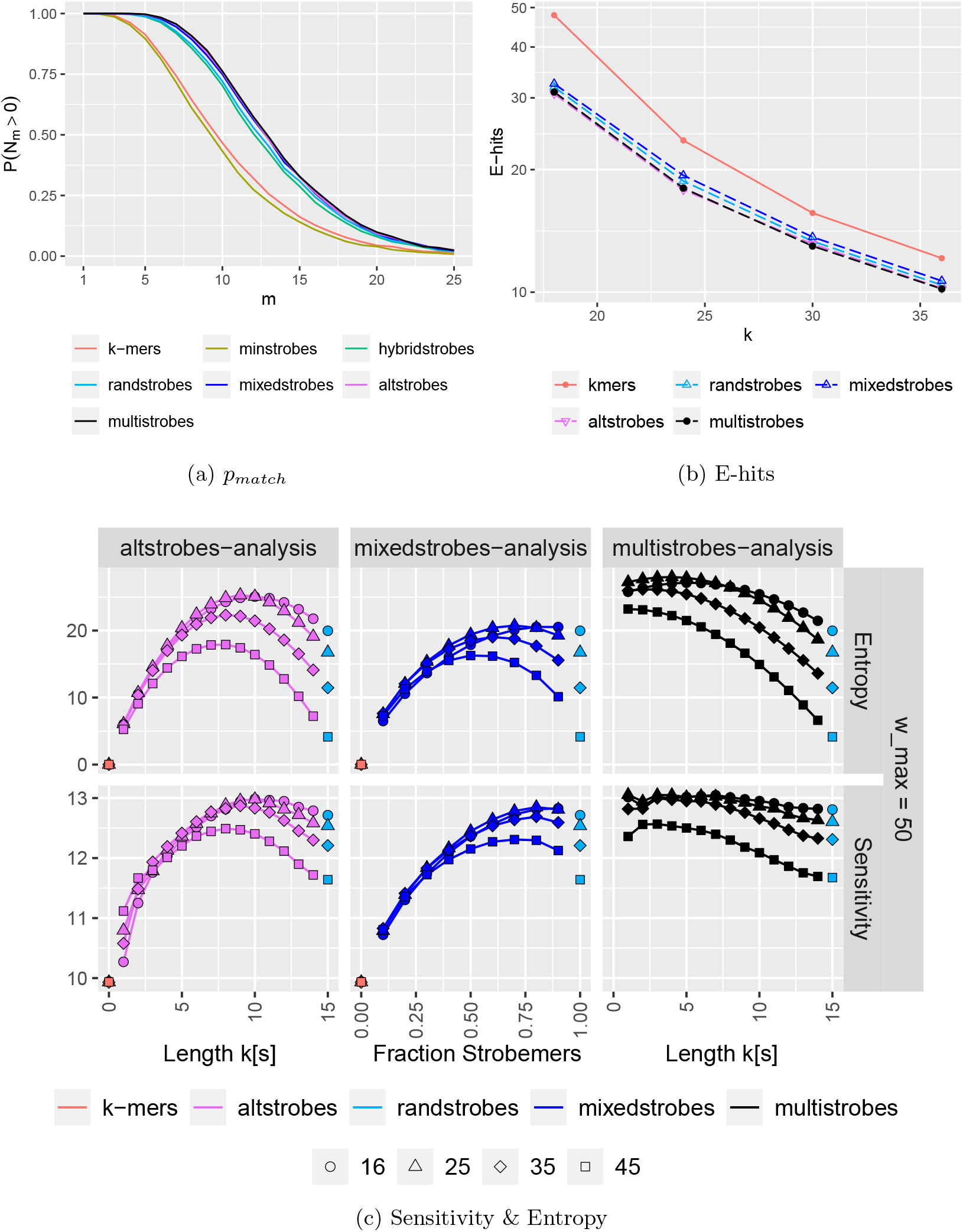
Simulations showing how pseudo-randomness in seed construct influence probability of *w* consecutive seeds producing at least one match in a region of length 2*w* = 128 between sequences. Panel A shows *P* (*N*_*m*_(30, 64) *>* 0) for seed constructs *k*-mers (*k*=30), minstrobes, hybridstrobes, and randstrobes with (2,15,25,50), mixedstrobes (2,15,25,50,0.8), altstrobes (2,10, 20,25,50), and multistrobes (2,5,25,25,50). Each *P* (*N*_*m*_(30, 64) *>* 0) estimate is derived from 10,000 instances of pairs of strings *S* and *T*. In general, a large gap is observed between non-random constructs (*k*-mers, minstrobes) to constructs with pseudo-randomness (hybridstrobes, randstrobes, mixedstrobes, altstrobes) for most mutation frequencies. The total sum of probabilities across *m* is higher for constructs with more random appearance. Panel B shows the seed uniqueness as expected number of hits (E-hits) from a seed randomly drawn from Human chromosome 21. Chromosome 21 of the human GRCh38 assembly was seeded with *k*-mers, randstrobes (2, *k/*2, 25, 50), mixedstrobes (2, *k/*2, 25, 50, 0.8), altstrobes (2, *k/*3, 2*k/*3, 25, 50) and multistrobes (2, 5, *k* − 5, 25, 50), whereby the number of extracted nucleotides (*k* = 30) was the same for all seeding techniques. Panel C (Entropy) shows *H*(**X**) for *k*-mers, randstrobes, altstrobes (for different (*k*_*s*_, *k*_*l*_), mixedstrobes (for different *q*), and multistrobes (for different *k*_*s*_). Panel C (Sensitivity) shows *P* (*N*_*m*_(30, 64) *>* 0) summed over *m* ∈ [1, 38], for various window sizes (*w*_*min*_, 50), (*w*_*min*_ ∈ 16, 25, 35, 45).

Next, we study how seed entropy is related to sensitivity at a more fine-grained level. Namely, we computed the seed entropy for *k*-mers, randstrobes as well as altstrobes, mixedstrobes, and multistrobes for different parameter settings varying (*k*_*s*_, *k*_*l*_), *q*, and evaluated them to empirical estimations of sensitivity summed over various error rates (simulation details in Suppl. section S3). Figure 2C (Entropy) shows entropy estimates for the constructs when using the same parameters as in [37], namely *k*=30 for *k*-mers, and (2, 15, 25, 50) for randstrobes which gives valid (30, 64)-seeds. We consequently set for mixedstrobes (2, 15, 25, 50, *q*) with *q* = 0, 0.1, …, 1.0, for altstrobes and multistrobes (2, *k*_*s*_, *k*_*l*_, 25, 50) with *k*_*s*_ = 1, 2, …, 14. Even though multistrobes and altstrobes are degenerate for *k*_*s*_ = 1 and 2 (hash values and high seed uniqueness cannot be guaranteed as discussed in section 2.6), we plot all values for all the parametrizations.

Overall, we see that our entropy model predicts predicts the sensitivity well (Fig. 2C) both between parameters within a seed construct and between seed constructs. Specifically, with our entropy model, we capture four trends. First, our model immediately suggests that in most cases, narrower (*w*_*min*_, *w*_*max*_) leads to lower entropy, hence, lower sensitivity. This result may seem obvious in hindsight, but it was unknown to us at the time of the strobemers study [37]. Second, given a fixed *w*_*min*_, we can typically predict which parameter settings of *q, k*_*s*_ yields good seed sensitivity for the constructs individually. The exception are mixedstrobes entropy peaks for *w*_*min*_ = 25, 35, and 45 which are slightly shifted to predict peak sensitivity at a lower *q* (0.5-0.9) than what is observed (0.7-0.9). This misprediction is more present with smaller windows such as *w*_*min*_ = 45. We are unable to explain this disagreement.

Third, we can compare *k*-mers, randstrobes, mixedstrobes, altstrobes, and multistrobes to each other. For most *w*_*min*_, we observe that multistrobes reach the highest peak entropy, followed in order by altstrobes, mixedstrobes, randstrobes, and finally *k*-mers. This trend is also present in the sensitivity curves. Ignoring the “degenerate” cases where *k*_*s*_ = 1 which we will discuss later, the peak sensitivity across all methods (13.0495) was reached by multistrobes with *k*_*s*_ = 4 closely followed by *k*_*s*_ = 3 (13.0494) for *w*_*min*_ = 25. After that, multistrobes with *k*_*s*_ = 7 for *w*_*min*_ = 16 (13.048). Several other *k*_*s*_ on the multistrobes curves for *w*_*min*_ = 16 and 25 also reached a summed sensitivity above 13. For altstrobes the peak sensitivity (12.980) was reached by *k*_*s*_ = 10 for both *w*_*min*_ = 16 and *w*_*min*_ = 25. For mixedstrobes the peak sensitivity (12.848) was reached by *q* = 0.8 and *w*_*min*_ = 25. These values roughly agrees with the peak entropies of the individual curves. Peak entropy is obtained by multistrobes with *k*_*s*_ = 4 (27.98), agreeing with peak sensitivity.

Fourth, the relative increase in entropy correlates well with the relative increase in sensitivity. For example, compare the relative distances between entropy and sensitivity peaks for *k*-mers, randstrobes, altstrobes, mixedstrobes, and multistrobes. Although the entropy curves are in general more spread out, the relative distances are relatively well preserved. In Suppl. Fig. S1, we can see that our observations generally also hold for experiments with *w*_*max*_ = 100 and 200 with varying *w*_*min*_. For example, lower window sizes typically perform worse than larger windows, and the parametrizations for a given construct and *w*_*min*_ can be ranked similarly as for the *w*_*max*_ = 50 experiments. However, we see a trend that with larger windows, both the sensitivity and entropy curves flatten out, suggesting that variable strobe sizes do not matter as much for sensitivity as for smaller windows. We believe this is because, in a large enough window, there are more possibilities to find error-free stretches longer than the strobe length, diminishing the necessity of hashing short strobes that can fit between mutations.

#### The degenerate case of *k*_*s*_ = 1

Interestingly we observed that multistrobes with *k*_*s*_ = 1 obtained the best sensitivity (13.054) across experiments for *w*_*min*_ = 25 and *w*_*max*_ = 50 (Fig. 2C). For other experiments, e.g., *w*_*min*_ = 25 and 50 with *w*_*max*_ = 100 (Suppl. Fig S1) the sensitivity deteriorated. With *k*_*s*_ = 1, the hash function can select at most four unique strobe length combinations according to the construct of multistrobes. Since the hash function in python use a different seed at each program start, we believe that if by chance a beneficial distribution is chosen (intuitively, *e*.*g*., *k*_*s*_ of 3, 7, 11, 15 to maximize length diversity) it may produce high sensitivity, but if a poor one is given (*e*.*g*., 1, 2, 3, 4, or collisions) it deteriorates results. This observation merits further studying the distribution of strobe lengths, as a uniform distribution over the range [*k*_*s*_, *k*_*l*_] may not be optimal.

#### Model limitations

During our study we learned that entropy is not the only feature that predicts seed sensitivity. An aspect not captured by our model is the probability that a contiguous segment (e.g., strobe or *k*-mer) is destroyed by mutations. An example of why modelling of this is needed is the following. Consider the following two different approaches of sampling mixedstrobes. In approach 1, we sample a *k*-mer or a strobemer based on the hash value of the first *k* nucleotides at the start of the seed. In approach 2, we sample a *k*-mer or a strobemer based on the hash value of the first *k/*2 nucleotides (the strobe length) at the start of the seed. It is straightforward to see that for any mutations *m >* 1, there will be more shared *k/*2-mers than *k*-mers between the sequences. Hence, the probability of sampling the same seed, and therefore generating a match, is higher for approach 2 (which we implement). The same argument holds for altstrobes and multistrobes, which is why we decide the strobe length based on the hash of *k*_*s*_.

Our model is agnostic to this probability. This becomes apparent when applying our model to strobemers with very narrow window sizes. We computed entropies and sensitivity estimates for mixedstrobes, altstrobes, and multistrobes with window sizes (*w*_*min*_,*w*_*max*_) of (49,50), (99,100), and (199,200) (Suppl. Fig. S2). In these cases, the seeds roughly acts as spaced *k*-mers but with randomness over strobe-size (altstrobes and multistrobes) or strobe fraction (mixedstrobes). While the entropy curves vary and predicts clear optima for these window sizes, the sensitivity curves are relatively flat but with peaks for *k*-mers, or near k-mer constructs (*k*_*s*_ = 1 altstrobes and mixedstrobes), which is expected when comparing k-mers to spaced k-mers when indels occur. Any indel within the seeds will destroy the seeds, and thus, give low sensitivity when indels are present at significant fractions. However, our model still estimates positive and variable entropy because the pseudo random selection of strobe lengths (for altstrobes and multistrobes) or seed type (for mixedstrobes). The model could therefore be improved by adding a probability distribution over indels, or that a contiguous region is error free.

Also, our model assumes a perfectly random hash function of linking. This is not true in practice, e.g., as we show in Suppl. Fig. S3. Also, implementation specific limitations to achieve uniform hashing for randstrobes has been identified [38]. Finally, as mentioned, our entropy measure cannot estimate entropy for pseudo-random seed constructs that are correlated between neighboring seeds such as minstrobes and hybridstrobes. However, our results show that the entropy of independent pseudo-random seed constructs as computed by our model overall predicts well the relative sensitivities of pseudo-random seed constructs.

### 3.2 Sequence matching results

We evaluated altstrobes (*k*_*s*_ = 10), multistrobes (*k*_*s*_ = 5) and mixedstrobes (*q* = 0.1, …, 1.0) against *k*-mers, and the other strobemers. We used *k*_*s*_ = 10 and *k*_*s*_ = 5 for, altstrobes and multistrobes, respectively, as according to our sensitivity analysis, they should be better performing than randstrobes for (*w*_*min*_, *w*_*max*_) = (25, 50). We used the sequence match analysis performed in [37], where several different aspects of sequence matching performance were evaluated, using both the simulations and genomic Oxford nanopore reads from [37]. For details about the data and simulation setup, see Suppl. Section S4. First, to verify our results from the sensitivity analysis, we performed an in detail analysis of altstrobes for different *k*_*s*_ using the match analysis on simulated data performed in [37] (see Suppl. Fig. S4). This analysis confirmed that *k*_*s*_ = 10 was preferable, in concordance with our sensitivity analysis. We then ran the matching analysis on all strobemer constructs. For comparability with the strobemer study [37], we used the same parameters (*e*.*g*., *k* = 30, and strobemers with (2, 15, 25, 50)). We included in the results minstrobes, hybridstrobes, and altstrobes seeded mixed with *k*-mers at different fractions *q* = 0, 0.1, …, 1.0. We also computed altstrobes with various *k*-mer fractions (i.e., mixed-altstrobes) for formatting consistency with the other results. Multi-strobes was not seeded mixed with *k*-mers, and therefore always appear at only fraction *q* = 1.0. We included results for strobemers of orders *n* = 2, 3 and 4, but we focus on evaluating *n* = 2 here.

Both the simulated data (Suppl. Fig. S5, panels with two strobes) and biological data (Fig. 3, Suppl. Fig. S6 and S7) experiments confirmed our sensitivity analysis. First, mixedstrobes with a strobemer fraction of roughly 70-80% perform better than strobemers-only seeding, also in this sequence matching analysis. The fraction of matches is higher for mixedstrobes at 80% than when seeding only randstrobes while, at the same time, the sequence coverage and expected island size were also better. Similar results can be observed for hybridstrobes and minstrobes. Secondly, altstrobes and multistrobes are outperforming randstrobes and mixedstrobes, whereby multistrobes have the most desired performance both on simulated and biological data, agreeing with our sensitivity analysis. Interestingly, for the biological data we observe that adding 20% *k*-mers to altstrobes further increases the sequence coverage over only using altstrobes. We believe this is because biological errors are less uniform, which may be beneficial for *k*-mers. As such mixed-altstrobes are using 10,20, and 30 as sampled strobe lengths, it could indicate that using a non-uniform distribution of strobe lengths in multistrobes (as discussed in the sensitivity analysis) could be beneficial.

**Fig. 3:**
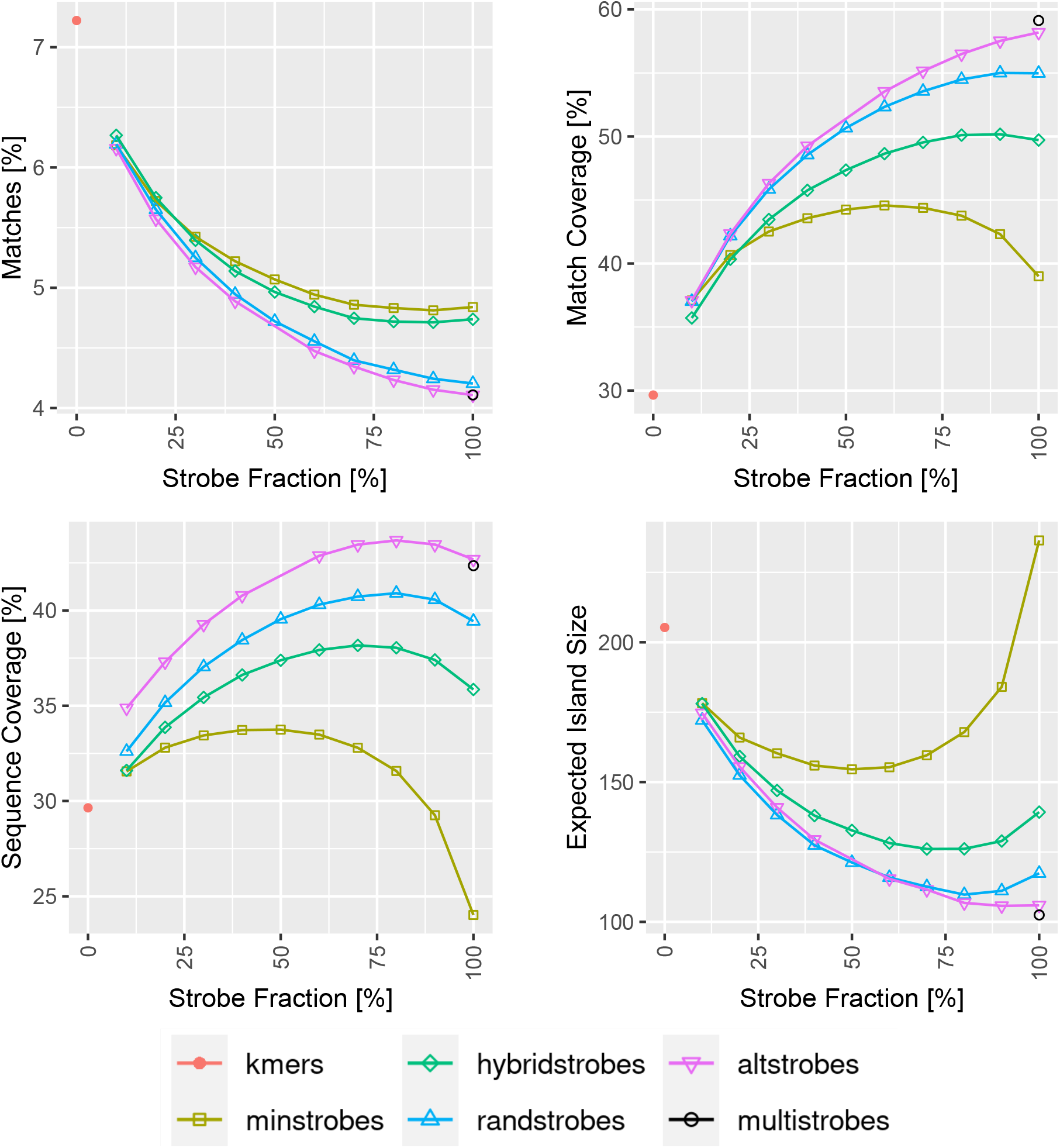
Comparison between (mixed-)strobemers (2,15,25,50, *q*), (mixed-)altstrobes (2,10,20,25,50, *q*), multistrobes (2,5,25,25,50) and *k*-mers (*k* = 30) when mapping genomic Oxford Nanopore Technology (ONT) reads from *E*.*coli* to its reference. The *E*.*coli* reads were split up in long disjoint segments of 2,000nt. Next, the segments were seeded with strobemer fractions *q* from 0% (*k*-mers) to 100% (strobemers), downstream windows set to [25,50] and all strobes combined adding up to equal length subsequences of size 30 for better comparison. Then for each segment, the collinear solution of raw hits was computed to subsequently quantify number of matches, match coverage, sequence coverage and expected island size.

### 3.3 Variable substitution frequency models

It is well documented that the frequency of nucleotide substitutions and insertions/deletions (indels) is species-specific and can vary across functional components of genomes. Hence, it is important to take these patterns and frequencies into account when benchmarking different seeding approaches. Previous study of strobemers [37] only investigated equally distributed substitutions, insertions, and deletions at probability 1/3. Here we include an analysis over various substitutions rates (from 0% to 100%) including also spaced *k*-mer seeds constructed as in [37]. Our analysis demonstrate that while spaced *k*-mers are the method of choice when analyzing sequences with low relative fractions of indels (0-10%), their performance deteriorate as indels become more prevalent (more than 10%) (Fig. 4). In contrast, the performance of strobemers and *k*-mers are more stable with varying indel rates.

**Fig. 4:**
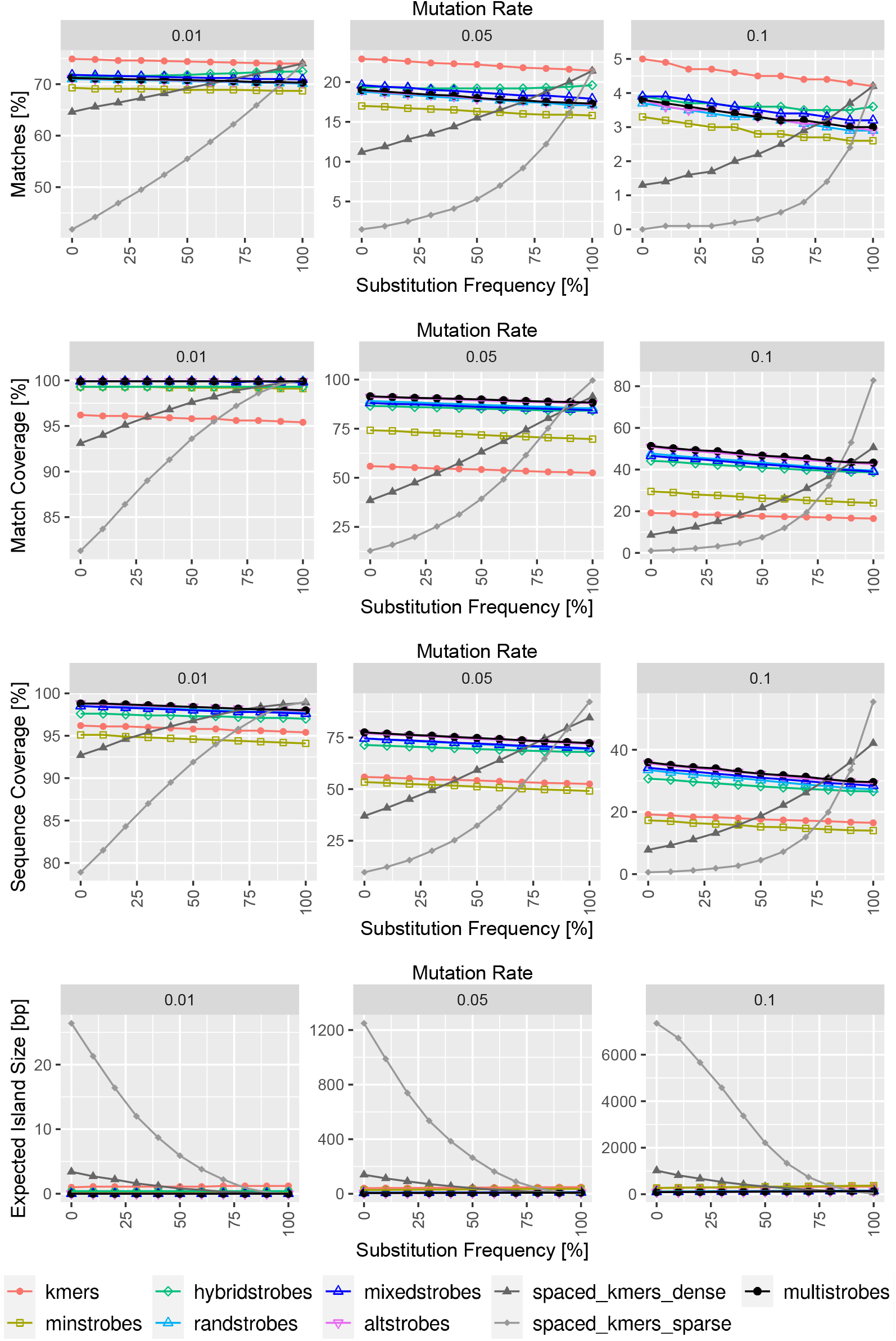
Performance of (spaced) *k*-mers and strobemers to various indel/substitution patterns in simulated data. For all the experiments, 1,000 random sequences of length 10,000nt were created and 1%, 5% and 10% of the nucleotides of the reference string were mutated for the different experimental conditions. Substitutions were hereby added with probabilities from 0% to 100% (substitution frequency), while the other mutation positions were filled with insertions and deletions with equal probability. Subsequently, the sequences were seeded with (spaced) *k*-mers of length *k* = 30 as well as strobemers of order *n* = 2 with all sub-strobes summing up to *k* = 30 and downstream windows set to [25,50]

### 3.4 Time and memory to construct altstrobes and mixedstrobes

We implemented altstrobes, mixedstrobes and multistrobes in StrobeMap [37] in C++ (Suppl. Fig. S8). *K*-mers are 3.5 times, 3.5 times, and 2.5 times faster than altstrobes, multistrobes and mixedstrobes, respectively. However, the time difference becomes negligible when looking at the total indexing time (including, *e*.*g*., sorting seeds and adding to hash table), especially when also taking into account that indexing is not the time limiting factor in most applications. The size on the index is nearly identical (Suppl. Fig. S8).

### 3.5 Minimap2 implementation

We implemented subsampled randstrobes, mixedstrobes, altstrobes and multistrobes in minimap2 [29] (see Suppl. section S5) to benchmark speed and accuracy of our seeding techniques and aligned simulated reads at various error rates to CHM13 (for details see Suppl. section S4). Our results indicate that altstrobes (2,9,18,25,50), mixedstrobes (2,14,25,50,0.8) and multistrobes (2,6,22,25,50) have slightly more (0.2%) correctly mapped reads, and slightly faster alignment time (up to 30%) compared to *k*-mers (*k* = 28) with a similar number of extracted seeds and peak RAM usage (Suppl. Fig. S9). Altstrobes, mixedstrobes and multistrobes also speed up alignment up to 3.5 times compared to default setting (*k* = 15). However, the number of correctly mapped reads remains lower than default setting. This is expected as using much smaller seeds is beneficial for sensitivity, at the cost of computing time. Also, minimap2’s search and extend parameters are highly optimized for exact *k*-mer seeds and minimizer subsampling, which is distorts the entropy of altstrobe, mixedstrobe and multistrobes seeds and is not something we modelled in our analysis.

### 3.6 Implementation and Software Availability

All the scripts used for the analysis and evaluation, as well as our seed implementations in StrobeMap and minimap2 are available at GitHub (https://github.com/benjamindominikmaier/mixedstrobes_altstrobes). Data analysis was conducted in R 4.1.3 [35] and Python 3.10 [46]. Figures were produced using the package ggplot2 3.3.5[48] and ggh4x 0.2.3[45] in tidyverse 1.3.1 [49] (R); and matplotlib 3.5.1 [23] and seaborn 0.11.2 [47] (Python).

## 4 Discussion

To our knowledge, we believe that we have provided the first study on seed sensitivity analysis for seeds that employ a pseudo-random sampling decision. We discovered a strong relationship between the pseudo-randomness of a seed construct and its effect on seed sensitivity, and we have experimentally verified that this relationship exists by modeling the entropy of a seed (Fig. 2, Suppl. Fig. S1). We discussed the cases where our model disagrees with our observations (for very short sampling windows; Suppl. Fig. S2) and explained why our model is incomplete. We have also expanded the strobemer family with mixedstrobes (combining *k*-mers and strobemers) and altstrobes (alternated strobe lengths), and multistrobes (generalizing altstrobes). We experimentally verified that for most parameter settings where they have higher entropy than randstrobes (the previously best performing strobemer), they also produce higher seed sensitivity. We further validated the benefit of using mixedstrobes, altstrobes, and multistrobes as seeds using several metrics from [37] (Fig. 3 and Suppl. Fig. S5) and also showed that altstrobes and multistrobes have lower repetitiveness than, e.g., randstrobes (Fig. 2B and Suppl. Fig. S10). Furthermore, we showed that mixedstrobes, altstrobes, and multistrobes are fast to construct (Suppl. Fig. S8-9) and do not constitute a bottleneck in mapping applications. Finally, we implemented randstrobes, mixedstrobes, and altstrobes in mininmap2 [29]. Minimap2 employs subsampling of seeds which distorts the relative entropies. Also, minimap2 implements chaining and other search-based cut-offs centered around minimizers. Nevertheless, we observed that using subsampled randstrobes and mixedstrobes within minimap2 for the most divergent sequence (10% mutation rates) both reduced runtime compared to *k*-mers of the same size with 25-30% and resulted in 0.2% more correctly mapped reads on CHM13 (Suppl. Fig. S9).

### 4.1 Future work

We believe that our work opens up for future work in several directions. Firstly, we may use our work’s insights to produce even better seed constructs. For example, our model suggests that finding seed constructs with higher entropy could improve sensitivity further. Another example is that, guided by our results (for *k*_*s*_ = 1; Fig. 2), it seems viable to investigate multistrobes with a different sampling distributions over sizes, such as only using a subset of strobe lengths, *e*.*g*., 3,7,11,15 or similar. Secondly, we believe that incorporating probabilities of error-free runs [3] will improve our model, which is currently only modeling entropy. Thirdly, it is common to subsample of seeds to reduce memory footprint and processing time. We are interested in adapting our model to incorporate subsampling. It is clear that when subsampling, the advantage that pseudo-random seed constructs (e.g., strobemers) have over *k*-mers reduces (Suppl. Table S1; also shown in [37]). This is because the high overlap of *k*-mers is removed with subsampling. Nevertheless, it would, for example, be beneficial to understand which subsampling densities and methods are suitable for pseudo-random seeds. Fourthly, since the minimap2 implementation is centered around minimizers, it is possible that aligners customized for, e.g., strobemers or other pseudo-random seeds may enjoy an even more substantial performance gain, as shown for short-read alignment [39].

## 5 Conclusion

Inspired by the many spaced seed studies on how to design optimal spaced seeds, we studied if the sensitivity of pseudo-random seeds can be predicted. We describe a model for estimating the entropy (randomness) of a seed, and we discover that seed sensitivity can be well predicted by the entropy. To our knowledge, we provide the first study that can predict seed performance for seeds that employ a pseudo-random sampling decision such as strobemers. Such seed have the advantage over, *e*.*g*., spaced *k*-mers in that they can match over indels. We discuss potential improvements to our entropy model as a predictor of sensitivity. Furthermore, we provide three new seed constructs, mixedstrobes, altstrobes, and multistrobes that are fast to construct and practical to use and can, for some parametrizations, improve over randstrobes, the previously most sensitive strobemer seed.

## 6 Acknowledgements

The computations were performed on resources provided by the Swedish National Infrastructure for Computing (SNIC) at Uppsala Multidisciplinary Center for Advanced Computational Science (UPPMAX) partially funded by the Swedish Research Council through grant agreement no. 2018-05973.

