## supplementary data for "Entropy predicts sensitivity of pseudo-random seeds"

#### Contents

**S.1 Construction of mixedstrobes and altstrobes** **SI-2**
S.1.1 Construction of mixedstrobes . . . . . SI-2
S.1.2 Construction of altstrobes . . . . . SI-2
S.1.2.1 Minimium $k_s$ for altstrobes . . . . . SI-5
S.1.2.2 Selecting altstrobes parametrization for experiments . . . . . SI-5
S.1.2.3 Altstrobes implementation details . . . . . SI-5
S.1.3 Construction of multistrobes . . . . . SI-5

**S.2 Example computations of $P(X_i|Y_j)$** **SI-8**
S.2.1 $k$-mers . . . . . SI-8
S.2.2 Randstrobes . . . . . SI-8
S.2.3 Altstrobes . . . . . SI-8
S.2.4 Mixedstrobes . . . . . SI-9
S.2.5 Multistrobes . . . . . SI-9

**S.3 Empirically estimating $P(N_m > 0)$** **SI-9**

**S.4 Experiments** **SI-9**
S.4.1 Evaluation metrics . . . . . SI-9
S.4.2 Data simulation . . . . . SI-10
S.4.3 Simulated results . . . . . SI-11
S.4.4 *E.coli* Oxford Nanopore Technology Reads . . . . . SI-11
S.4.5 Biological Data . . . . . SI-12

**S.5 Minimap2 implementation details** **SI-12**
S.5.1 Minimap2 analysis . . . . . SI-12

**S.6 Figures and tables** **SI-12**

### S.1 Construction of mixedstrobes and altstrobes

#### S.1.1 Construction of mixedstrobes

Mixedstrobes consists out of a specified fraction of $k$-mers and strobemers and may be sampled with either of the three strobemers seeding methods (minstrobes, hybridstrobes and randstrobes), but we will only consider randstrobes here.

Whether a strobemer or a $k$-mer is seeded depends on the hash value of the first strobe $h(S[i : i + \ell])$ and the user-defined strobe fraction $q$. For instance, for mixedstrobes with $q = 0.8 = 80\%$, strobemers should be generated when $h(S[i : i + \ell)$ % $100 < 80$; otherwise a $k$-mer should be sampled instead. Strobemers are sampled following the routine [SR2]. When sampling a $k$-mer $(1 - q)$, $n\ell$ consecutive nucleotides are taken starting from the start position of the first strobe ($S[i : i+n\ell]$) and converted to its respective hash value (Python). By taking $n\ell$ nucleotides, we obtain the same subsequence lengths as strobemers consisting out of $n$ strobes of length $\ell$. In C++, $k$-mers are constructed by summing the hash values of $S[i + j * \ell]$, which are each divided by $(n + j)$ to guarantee non-commutativity, whereby $j \in [1, n]$.

#### S.1.2 Construction of altstrobes

Altstrobes are modified randstrobes where the strobe length is alternating between shorter ($k_s$) and longer strobes ($k_l$) with $|k_s| + |k_l| = k$. Whether the first strobe is of length $|k_s|$ or $|k_l|$ is decided based on the hash value of the substring of length $|k_s|$ (*i.e. the potential first strobe).* Note that it is highly advised against making the decision based on the hash value of $k_l$ as this may lead to unnecessarily many seeds being destroyed where ($k_s$, $k_l$)-altstrobes should have been sampled and there is are mutations within the positions $[k_s, k_l]$ downstream from the start position of the seed. Furthermore, one should not seed altstrobes where the length of the shorter strobe is smaller than 5 to avoid seed repetitiveness (low uniqueness).

In case, that we are sampling mixed-altstrobes, the hash value is divided by 100, whereby the fractional part (remainder) is discarded. This integer division is required as we have to make two independent decisions using the hash value: whether to sample an altstrobe or a $k$-mer and in case of an altstobes whether to sample a short or a long strobe first.

The following downstream strobes are selected from a window W by alternatively sampling a short and a long strobe using the randstrobe linking routine as described in [SR2]. We adjust the offset of $(w_{min}, w_{max})$ depending on if it is the long or short strobe we sample. Specifically, we let $k_l$ in altstrobe $(k_s, k_l)$ be sampled from $[w_{min} - (k_l - k_s)/2, w_{max} - (k_l - k_s)/2]$ and $k_s$ in altstrobe $(k_l, k_s)$ be sampled from $[w_{min} + (k_l - k_s)/2, w_{max} + (k_l - k_s)/2]$. This guarantees that the maximum length of the altstrobe seed remains the same as randstrobes ($w = w_{max} + k/2$) which is important for benchmarking.

In the default altstrobe protocol where $2k_s = k_l$, we store $S[i]$ for short strobes and $S[i]$ as well as $S[i] + k_s$ for long strobes. This allows us to keep track of whether $(|k_s|, |k_l|)$ or $(|k_l|, |k_s|)$ was sampled and facilitates analysis (e.g. sequence coverage) as now all strobes are of equal length ($k_s$). If generalized altstrobes are sampled where $2k_s \neq k_l$ and information about the exact strobe composition is required, we store all strobe positions.

**Algorithm 1** Mixedstrobes

**Function:** Mixedstrobes $(S, n, \ell, w_{min}, w_{max}, s)$
**Input:** Sequence $S$, number of strobes $n$, strobe lengths $\ell$, strobe window $W[w_{min}, w_{max}]$, strobe fraction $s$
**Output:** Mixedstrobes of order $n$ and their positions from $S$

$s = \frac{N}{D}$ `// Split strobe fraction` $s$ `in numerator` $N$ `and denominator` $D$
$O = []$ `// Initialize array of strobemers and their positions`
**for** $i \in [1, |S| - (n * \ell + 1)]$ **do** `// Iterate over all positions`
    $m_1 = S[i : i + \ell]$
    $P = [i,]$
    $H = h(m_1)$
    **if** $h(m_1)\ \%\ D < N$ **then** `// Sample strobemer at position` $i$
        $w_u = min(w_{max}, ([S] - i)/(n - 1))$ `// Second argument only active at end of S`
        $w_l = max(w_{min} - (w_{max} - w_u), \ell)$
        **for** $j \in [2, n]$ **do**
            $w' = [i + w_l + (j-2)w_u, i + (j-1)w_u]$ `// Window to look for current strobe`
            $p = \arg\min_p \{p : h(m \bigoplus S[p : p + \ell]) \leqslant h(m \bigoplus S[i' : i' + \ell]), \forall i' \in w'\}$ `// Selecting strobes based on strobemer protocol, here exemplified using randstrobes)`
            $P \mathrel{+}= p$
            $H \mathrel{+}= (-1)^j * j * h(S[p : p + \ell])$ `// Non-commutative hash value combination`
        **end**
    **else** `// Sample` $k$`-mer of length` $n\ell$ `at position` $i$
        **for** $j \in [2, n]$ **do**
            $P \mathrel{+}= i + j * \ell$
            $H = h(S[i : i + n\ell])$
        **end**
    **end**
    $O \mathrel{+}= (P, H)$
    **return** $O$
**end**

**Algorithm 2** Altstrobes

```
Function: Altstrobes (S, n, k_s, k_l, w_min, w_max)
Input: Sequence S, number of strobes n, length of short and long strobes k_s and k_l,
       strobe window W[w_min, w_max]
Output: Altstrobes of order n and their positions from S
Require : n ∈ 2ℤ
O = [] // Initialize array of altstrobes and their positions
for i ∈ [1, |S| − (n ∗ ℓ + 1)] do // Iterate over all positions
    P = [i,]
    if h(S[i : i + k_s]) % 2 = 0 then // Sample altstrobe(k_s,k_l) at position i
        H = h(S[i : i + k_s])
        strobes = [k_s, k_l]
    else // Sample altstrobe(k_l,k_s) at position i
        H = h(S[i : i + k_l])
        strobes = [k_l, k_s]
    end
    w_u = min(w_max, ([S] − i)/(n − 1)) // Second argument only active at end of S
    w_l = max(w_min − (w_max − w_u), k_l)
    for j ∈ [2, n] do
        k' = strobes[(j + 1)%2] // Retrieve length of next strobe
        offset = (k_s + k_l)/2 − k'// Adjusting window offsets
        w' = [i + w_l + (j − 2)w_u + offset, i + (j − 1)w_u + offset] // Window to look for current strobe
        p = argmin_p{p : h(m ⊕ S[p : p + k']) ⩽ h(m ⊕ S[i' : i' + k']), ∀i' ∈ w'} // Selecting strobes based on randstrobe protocol
        P += p
        H += (−1)^j ∗ j ∗ h(S[p : p+k']) // Non-commutative hash value combination
    end
    O += (P, H)
    return O
end
```

###### S.1.2.1 Minimium $k_s$ for altstrobes

The combination of strobe lengths $k_s$ and $k_l$ matter in practice. A too short $k_s$ leads to degenerate seed constructs for two reasons. Firstly, the number of possible hash values is limited for very short strobes (e.g., 4, 16, and 64 hash values for 1-, 2-, and 3-mers respectively). Thus, it cannot be guaranteed that an even fraction of ($k_s$, $k_l$) and ($k_l$, $k_s$) is computed for such parametrizations, which decreases randomness in seed selection. Secondly, $|k_s|$ needs to be long enough to avoid random repetitions in substrings of length $w_{max} - w_{min}$. To see why this is the case, consider using a short strobe size of 2. If we have a sequence ACGACTACA..., where AC hash to an even number (short strobe selected), we will very likely link AC (short strobe) with the same $k_l$ multiple times as the downstream selection window of size $w_{max} - w_{min}$ will share many strobes for reasonably large windows. This deteriorates both randomness and uniqueness of seeds. We show uniqueness per $k_s$ length in Suppl. Fig. S3, where for random simulated sequences has a window size of 25nt. For this window size, $|k_s| \geq 6$ is needed to be competitive with other parametrizations in terms of uniqueness.

###### S.1.2.2 Selecting altstrobes parametrization for experiments

Several combinations of $(k_s, k_l)$ in altstrobes could lead to lower correlation and potentially outperform randstrobes, as was shown in Figure S4. We performed simulations on random sequences for all possible altstrobe combinations of combined strobe length of 30. We found that sequence coverage, match coverage and expected island size was best for altstrobes with $k_s$ between 7 and 10 (Fig. S4). Since simulated sequences are less repetitive than biological sequences, we opt for the longest possible $k_s$ with good metrics.

###### S.1.2.3 Altstrobes implementation details

Despite altstrobes orders being multiples of 2, we output the results as multiples of order 3 to keep track on whether a short-long or a long-short combination was sampled. As the long strobe $k_l$ is exactly double the size of $k_s$ we can store altstrobe seeds of order 2 as $(x_1, x_1 + k_s, x_2)$ and $(x_1, x_2, x_2 + k_s)$ with $x_1$ and $x_2$ being the start positions of the strobes and $k_s$ being the shorter strobe length.

##### S.1.3 Construction of multistrobes

Multistrobes are generalized altstrobes which allows to sample a range of possible strobe lengths ranging from $k_s$ to $k_l$ with $|k_s| + |k_l| = k$ within the same seeding pass.

As the strobe length combination is decided based on the hash value of a subsequence of length $k_s$ ($h(S[i : i + |k_s|])$), we have to deal with uniformity issues. For instance, for $|k_s| = 2$ and $|k_l| = 28$, there are only $4^2 = 16$ hash values $h(S[i : i + |k_s|])$ making it impossible to select all 27 options from $|k_l - k_s + 1|$. However, there are less strobe length combinations from $[k_s,\ k/2]$ than the 16 hash values. Therefore, if we first pick a strobe size from $[k_s,\ k/2]$ (14 options), we may in a second step choose whether it should be the first or the second strobe. In more detail, the approach is performed as follows.

First, a strobe length $k'$ is sampled from $[k_s,\ k/2]$ based on the hash value of $h(S[i : i + |k_s|]$ as

$$k' \doteq |k_s| + \big(h(S[i : i + |k_s|])\%(|k_l| - |k|/2 + 1)\big).$$

Second, we decide if the strobe of length $k'$ should come first or second based on

$$f(i,k,w,S,*) = \begin{cases} \text{Sample multistrobe } (k', k-k') & \text{if } \big((h(S[i:i+|k'|])\ //100)\%\ 2 = 0 \\ \text{Sample multistrobe } (k-k', k'), & \text{otherwise} \end{cases}$$

This approach can, at best, double the possible hash value selection options and also improve the uniformity of the number of short-long and long-short combinations when $k_s$ is small. Note however that our two-step implementation described above is not fully uniform when $|k_l - k_s + 1|$ is uneven, as the option $(k/2, k/2)$ is appearing with double probability as their short-long and long-short combinations are identical. The python implementation implements this two-step procedure to cope with very small $k_s$ for our sensitivity benchmarks while the C++ implements the sampling as described in the main paper for runtime.

**Algorithm 3** Multistrobes

**Function:** Multistrobes $(S, n, k_s, k_l, w_{min}, w_{max})$
**Input:** Sequence $S$, number of strobes $n$, minimum strobe length $k_s$, maximum strobe length $k_l$, strobe window $W[w_{min}, w_{max}]$
**Output:** Multistrobes of order $n$ and their positions from $S$
**Require :** $n \in 2\mathbb{Z}$

```
O = [] // Initialize array of multistrobes and their positions
for i ∈ [1, |S| − (n ∗ ℓ + 1)] do // Iterate over all positions
    P = [i, ]
    |k'| ≐ |k_s| + (h(S[i : i + |k_s|])%(|k_l| − |k|/2 + 1))
    if ((h(S[i:i+—k'—]) // 100) % 2 = 0) then // Decide whether to start with a short strobe
        k_1 = k'
        k_2 = k − k'
    else
        k_1 = k − k'
        k_2 = k'
    end
    strobes = [k_1, k_2]
    H = h(S[i : i + k_1])
    w_u = min(w_max, ([S] − i)/(n − 1)) // Second argument only active at end of S
    w_l = max(w_min − (w_max − w_u), k_2)
    for j ∈ [2, n] do
        k' = strobes[(j + 1)%2] // Retrieve length of next strobe
        offset = k/2 − k'// Adjusting window offsets
        w' = [i + w_l + (j − 2)w_u + offset, i + (j − 1)w_u] + offset // Window to look for current strobe
        p = arg min_p{p : h(m ⊕ S[p : p + k']) ⩽ h(m ⊕ S[i' : i' + k']), ∀i' ∈ w'} // Selecting strobes based on randstrobe protocol
        P += p
        H += (−1)^j ∗ j ∗ h(S[p : p+k') // Non-commutative hash value combination
    end
    O += (P, H)
    return O
end
```

#### S.2 Example computations of $P(X_i|Y_j)$

For convenience, we will denote $P(X_i|Y_j)$ for $k$-mers, randstrobes, mixedstrobes, altstrobes, and multistrobes as $P_k$, $P_r$, $P_{mi}$, $P_a$, $P_{mu}$, respectively.

##### S.2.1 $k$-mers

We have

$$P_k = \begin{cases} 1 & \text{if } i+j+2 \leq k \\ 0 & \text{otherwise.} \end{cases} \tag{1}$$

We will refer to this equation as $E_6(k)$ taking the length of the $k$-mer as argument, as it turns out handy to reuse for some of the other constructs.

##### S.2.2 Randstrobes

Let the strobe size be $k' = \lfloor k/2 \rfloor$. If $Y_j$ is located on the second strobe, the probability is $E_6(k')$ since $i$ can only be covered by the second strobe. If $j$ is located on the first strobe, $i$ can be either covered by the first or second strobe. Hence, we need to structure the probability up into cases. The possible second strobe coverings of position $i$ given $Y_j$ depends on the downstream window location, and is further restricted by the window size ($B = w_{max} - w_{min} + 1$) and the strobe length. The second strobe coverings is computed as

$$A = \min(k', B, i+j+2-w_{min}, w-(i+j+2)).$$

Under the assumption that $w_{min} > k'$, we have

$$E_7(k') = \begin{cases} 1 & \text{if } i+j+2 \leq k' \\ A/B & \text{if } w_{min} - j \leq i \leq w-(j+1) \\ 0 & \text{otherwise.} \end{cases} \tag{2}$$

Let $I_j$ be an indicator variable with $I_j = 1$ if $j$ is placed on first strobe and 0 otherwise. Then,

$$P_r = E_7(k')I_j + E_6(k')(1-I_j)$$

##### S.2.3 Altstrobes

For altstrobes, we have the same cases as for randstrobes ($E_6$ and $E_7$), but we need to sum over the scenario that either the first or the second strobe is short (with equal probabilities 0.5). We have

$$P_a = (0.5E_7(k_s) + 0.5E_7(k_l))I_j + (0.5E_6(k_l) + 0.5E_6(k_s))(1-I_j).$$

To guarantee the same maximum seed length we employed a window offset based on if the first strobe as short or long (as described in section 2.8), this window adjustment is computed before $E_7$ is computed.

##### S.2.4 Mixedstrobes

For mixedstrobes, we need to condition on the probability $q$ that either a randstrobe or a $k$-mer is sampled. We have

$$P_{mi} = (qE_7(k') + (1-q)E_6(k))I_j + qE_6(k')(1-I_j).$$

Where we have included in the event $I_j = 1$ that a $k$-mer is sampled, as it can be seen as a strobe of length $k$ with the second strobe being of length 0.

##### S.2.5 Multistrobes

Finally, multistrobes samples a strobe length $\ell \in [k_b, k-k_b]$ and sets $k-\ell$ as length for the second strobe. Let $p_\ell = \frac{1}{k-2k_b+1}$ denote this probability (which is uniform over the considered strobe lengths). Let the shorter of the two be denoted by $k_s$ and the longer be denoted by $k_l$, similarly to altstrobes. Then the probability is similar to $P_a$ but summed over possible strobe lengths. We have

$$P_{mu} = \sum_{\ell=k_b}^{k-k_b} p_\ell \Big( (0.5E_7(k_s) + 0.5E_7(k_l))I_j + (0.5E_6(k_l) + 0.5E_6(k_s))(1-I_j) \Big).$$

In case of $P_r$, $P_{mi}$, $P_a$, and $P_{mu}$, the above probabilities assume that the second strobe is chosen uniformly at random over the window, meaning that they assume a perfect random hash function. When the first strobe in altstrobes or multistrobes is very short it is not possible to sample uniformly from the window, as was discussed and shown in the construction of altstrobes section.

#### S.3 Empirically estimating $P(N_m > 0)$

We empirically estimated $P(N_m > 0)$ for each of the seed construct parametrizations as follows. A string $S$ (letters A, C, G, and T) of length $4w$ is simulated at random, and a second string $T$ is simulated by copying $S$ and randomly performing exactly $m$ mutations within the first $2w$ nucleotides of $S$, creating a fixed mutation rate within the segment of $2w$ on $S$. As for the mutation profile, we uniformly draw the substitution rate $s \in [0,1]$ and set insertions and deletions each to $(1-s)/2$. We then construct seeds from $S$ and $T$ with each seed construct and parametrization. We store the event that seeds from the first $w$ seeds on $S$ and $T$ has at least one match. We estimate $P(N_m > 0)$ from the fraction of experiments with at least one match out of $U$ experiment replicates, where $U = 10,000$ for $w_{max} = 50$ and $U = 1,000$ for $w_{max} = 100$ and 200. Finally, we obtain the summed sensitivity $\sum_{m=1}^{M} P(N_m > 0)$ to capture the sensitivity over a range of error rates. $M$ is chosen such that it corresponds to 30% error rate within the segment of length $2w$. We chose 30% as all seed constructs returned no matches with the given seed lengths for this error rate. We chose $w_{max}$ of 50, 100, and 200, giving rise to $w$ of 64, 114, and 214 respectively. We chose these values to study the effect of the window size, where 50 and 100 are also consistent with previous study [SR2].

#### S.4 Experiments

##### S.4.1 Evaluation metrics

We also evaluated our best performing seed constructs from the sensitivity analysis in a more general sequence matching scenario using previously designed metrics in [SR2], namely fraction of matches, match coverage, sequence coverage, and expected island size:

- **Fraction of Matches:** proportion of the query seeds that matched the reference
- **Match Coverage:** proportion of nucleotides covered by the $k$-mers and strobemers from end-to-end including potential gaps
- **Sequence Coverage:** proportion of nucleotides covered by the strobes of matches, so it distinguished from match coverage by disregarding the gaps between the strobes
- **Expected Island Size:** An island is the maximal interval of consecutive nucleotides without matches. If a random location from the reference genome is selected, they may either be covered by matches (size of island = 0) or islands of various length. For a sequence $S$ and a set of islands length $X$, the expected island size $E$ is computed as follows:

$$E = \frac{1}{\mid S \mid} \sum_{x \epsilon X} x^2 \tag{3}$$

.

We used the same parameters and simulation setup as in [SR2] (details in Suppl. Section S2). Our seed sensitivity measure investigated in section 3.1 is most related to the metrics sequence coverage and island E-size, which measures two important metrics for sequence matching. While some sequence comparison algorithms may require a high fraction of matches for accurate similarity estimation and therefore optimize for number of matches, it is not typically needed for, e.g., read mapping applications. For example, it was shown in [SR2] that $k$-mers produce the highest fraction of matches under a random error distribution, but the high fraction of matches often occur because of several consecutive overlapping matches. In [SR1], the authors argued that this is bad as many of the matches are redundant, and they aim to select combinations of seeds that yield matches which overlap as little as possible which does not optimize for a high fraction of matches.

##### S.4.2 Data simulation

The performance of mixedstrobes, altstrobes and multistrobes was benchmarked and compared to (spaced) $k$-mers and strobemers on simulated sequencing data in a similar scenario to the match evaluation in [SR2], described here for convenience. Random reference sequences of 10,000 nucleotides were simulated, whereby the probability of each of the four nucleotides was 25% for each position. To create the corresponding query sequence, 1%, 5% and 10% of the nucleotides of the reference string were mutated for the different experimental conditions. Insertions, deletions and substitutions were hereby added with equal probability of 1/3. To reduce sample variation bias, these simulations were repeated 1,000 times. The randomly-simulated references and queries were now used as inputs to seed $k$-mers, spaced $k$-mers, randstrobes, minstrobes, hybridstrobes, mixedstrobes, and altstrobes. All $k$-mer and strobemer parameters were set as in [SR2], namely $k = 30$ and (2,15,25,50), altstrobes with (2,10,20,25,50) and multistrobes with (2,5,25,25,50), all yielding valid $(30, 64)$-seeds. Then, mixedstrobes was sampled with (2,15,25,50,$q$), $q \in [0, 0.1, 0.2, \ldots, 1.0]$. We sampled mixedstrobes by combining each strobemer type (randstrobes, minstrobes, hybridstrobes, altstrobes) with $k$-mers as it was easy with our experiment setup.

##### S.4.3 Simulated results

Our results demonstrate that $k$-mers performed best regarding the fraction of matches (Fig. S5A), which decreased linearly with increasing fractions of strobemers. Contrary, the match coverage (Fig. S5B) and the sequence coverage (Fig. S5C) increased with higher strobemer fractions, even though it leveled off towards the highest strobemer fractions. Sequence and match coverage were shown to be best for multistrobes and pure altstrobes (100%) followed by randstrobes and hybridstrobes with a strobemer content of 70-80% (Fig. S5C), largely outperforming $k$-mers and even being slightly better than the pure strobemer implementations. We observed similar results when looking at the expected island size which displayed an exponential decay behavior when adding more strobemers, whereby the benefit of going beyond 80% strobemers was either non-existent or very low (Fig. S5D).

Overall, mixedrandstrobes with a strobe fraction of 70-80% were found to be superior to the pure strobemer implementations as higher fraction of matches and a lower island size was observed while maintaining high sequence and match coverage. Furthermore our analysis suggests that altstrobes are strictly better than the currently best known strobemer construct (randstrobes) as the fraction of matches, sequence coverage and match coverage are higher while the number of expected island sizes remained lower. The generalized implementation of altstrobes, multistrobes, performed even better as it is outperforming altstrobes in all matching metrics (Supp. Table 1).

As the spaced $k$-mer protocol performed worse than $k$-mers on all metrics (fewer matches, lower match coverage, and larger expected island size), their results are not displayed for better visualisation. However, full data including spaced $k$-mers is presented in Suppl. Table 1.

We also looked at strobemers of orders 2 to 5. We observe that in practice, $n \geq 4$ does not offer benefit over $n = 2$ and 3 for the application of strobemers that we consider in this study (Fig S5). Note that $k = 30$, can be divided by 2,3, and 5 without leaving a remainder, guaranteeing same number of positions sampled. Also, altstrobes and multistrobes were only computed for $n = 2, 4$ to guarantee same number of positions sampled as the other seeds. For strobemers of order 4, we observe that altstrobes and multistrobes perform very similar which can be explained by their shortest possible strobes being of identical length ($k_s = 5$) and the considerably lower number of strobe combinations (6 vs. 21) for multistrobes of order 4.

##### S.4.4 *E.coli* Oxford Nanopore Technology Reads

In contrast to simulated data where we compare query sequences and their corresponding references individually, biological data may contain spurious matches, which requires measurements of the sequence matching metrics on biological data to reaffirm our previous simulation results.

To this end, the one thousand *E.coli* Oxford Nanopore Technology reads used in [SR2] ranging from 17,360nt to 52,197nt (median 19,601nt) were split up in disjoint segments of 2,000bp before computing the collinear chain solution of the raw hits for each of the segments. The collinear chaining algorithm determines chains of matches that are in identical order in both sequences. Hence, the collinear solution can be viewed as a proxy for finding the true location of the hits by taking only the longest collinear chain of the hits into account, as in [SR2], to avoid overcounting "spurious" hits caused by local matches in repetitive regions that occur throughout the genome. Raw unmerged hits were assessed rather than non-overlapping approximately matchings (NAMs) to make the analysis identical to the simulated experiment. Subsequently, for each read, the number of matches, the match coverage, the sequence coverage and the expected island size were computed for the collinear solution of hits (Fig. 3 and Suppl.

Figs. S6 and S7).

##### S.4.5 Biological Data

The thousand *E.coli* reads were downloaded from Sequence Read Archive with Run ID SRR13893500 (available here: `https://trace.ncbi.nlm.nih.gov/Traces/?view=run_browser&acc=SRR13893500&display=download`. We selected the 1,000 longest reads that aligned to the *E.coli* genome with more than 95% of the total read length as in [SR2]. The reads were mapped to the *E.coli* genome assembly GCA 003018135.1 ASM301813v1 (available here: `https://www.ncbi.nlm.nih.gov/genome/167?genome_assembly_id=368373`). For the E-hits and uniqueness analysis we used human chromosome 21 (NC_000021.9) from GRCh38.p13 Primary Assembly. For the human data analysis with minimap2, we used assembly CHM13 v2.2 (available here: `https://s3-us-west-2.amazonaws.com/human-pangenomics/T2T/CHM13/assemblies/analysis_set/chm13v2.0.fa.gz`).

#### S.5 Minimap2 implementation details

We implemented randstrobes, mixedstrobes, altstrobes and multistrobes in the fast sequence mapping and alignment program minimap2. First, we find minimizer $k$-mers based on the shorter strobe $k_s$ (altstrobes and multistrobes) or the first strobe $k/2$ (randstrobes and mixedstrobes). To decide whether a $k$-mer or a randstrobe (mixedstrobes) should be sampled, a modulo operation is performed on the hash value of the first strobe (see mixedstrobe implementation). Analogously, a decision about $k_s$ and $k_l$, and $k_1$ and $k_2$ is made for altstrobes and multistrobes, respectively (see Section 2.6 of the paper).

Next, the second strobes are selected from a downstream window [25,50] that minimizes the function $\arg\min_p\{p : h(m \bigoplus S[p : p+\ell]) \leqslant h(m \bigoplus S[i' : i'+\ell]), \forall i' \in w'\}$. The hash values of the selected strobes are combined using non-commutative concatenation (hash values of linked $k$-mers) and are assigned to our fuzzy seeds of length $w_{max} + k - 1$. As it is not possible to sample seeds of these length from the reverse strand at the beginning of the sequence, only forward seeds are sampled for the first $w_{max} + k - 1$ nucleotides, and *vice versa* only reverse strand seeds at the end of the sequence.

##### S.5.1 Minimap2 analysis

To measure speed and accuracy of our newly implemented seeding methods, we benchmarked them against $k$-mers with $k = 15$ (default setting) and $k = 28$. To this end, we sampled 100,000 reads of length 10,000nt from random positions across the *Homo sapiens* (human) genome assembly CHM13 that did not contain any incompletely specified bases. Next, we inserted, deleted, or substituted nucleotides with equal probability of 1/3 each across the reads with the mutation rate $0.01, 0.05, 0.1$, as in [SR2], and converted the read sequence into its reverse-complement counterpart with probability of 1/2. These mutated reads were mapped and aligned back to the CHM13 assembly using minimap2 with default mapping and aligning settings. A read was considered to be mapped correctly if at least 1nt was mapped to its correct location.

#### S.6 Figures and tables

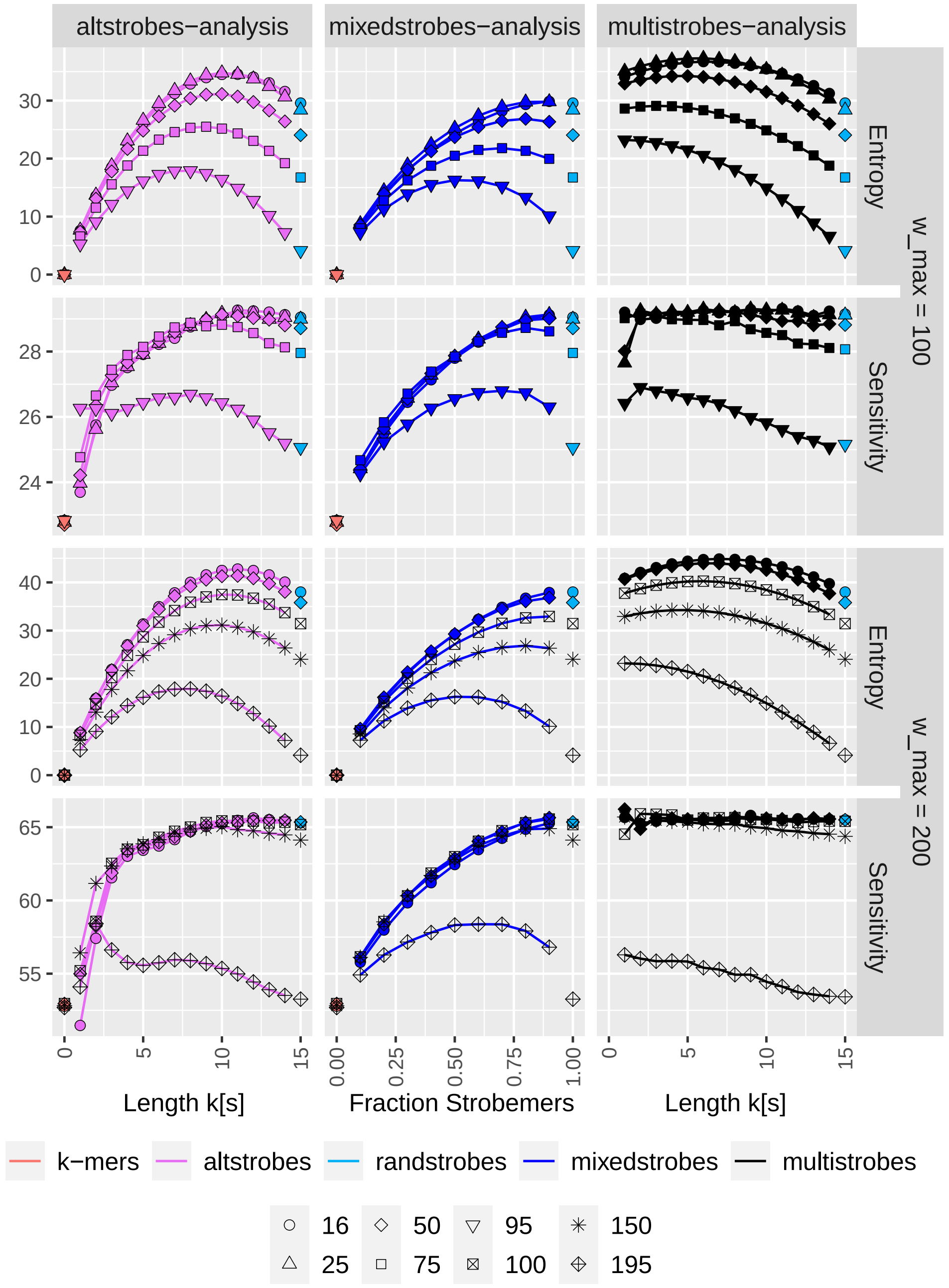


**Figure S1. Simulations showing how stochasticity in seed construct influence probability of $w$ consecutive seeds producing at least one match in a region of length $2w$=128 between sequences.**
Panel Entropy shows $H(\mathbf{X})$ for $k$-mers, randstrobes, altstrobes (for different $(k_s, k_l)$), mixedstrobes (for different $q$) and multistrobes (for different $(k_s, k_l)$). Panel Sensitivity shows $P(N_m(30, 64) > 0)$ summed over $m \in [1, 60]$ for various window sizes $(w_{min}, 100)$, $(w_{min} \in 16, 25, 50, 75, 95)$ and $(w_{min}, 200)$, $(w_{min} \in 16, 50, 150, 190, 195)$.

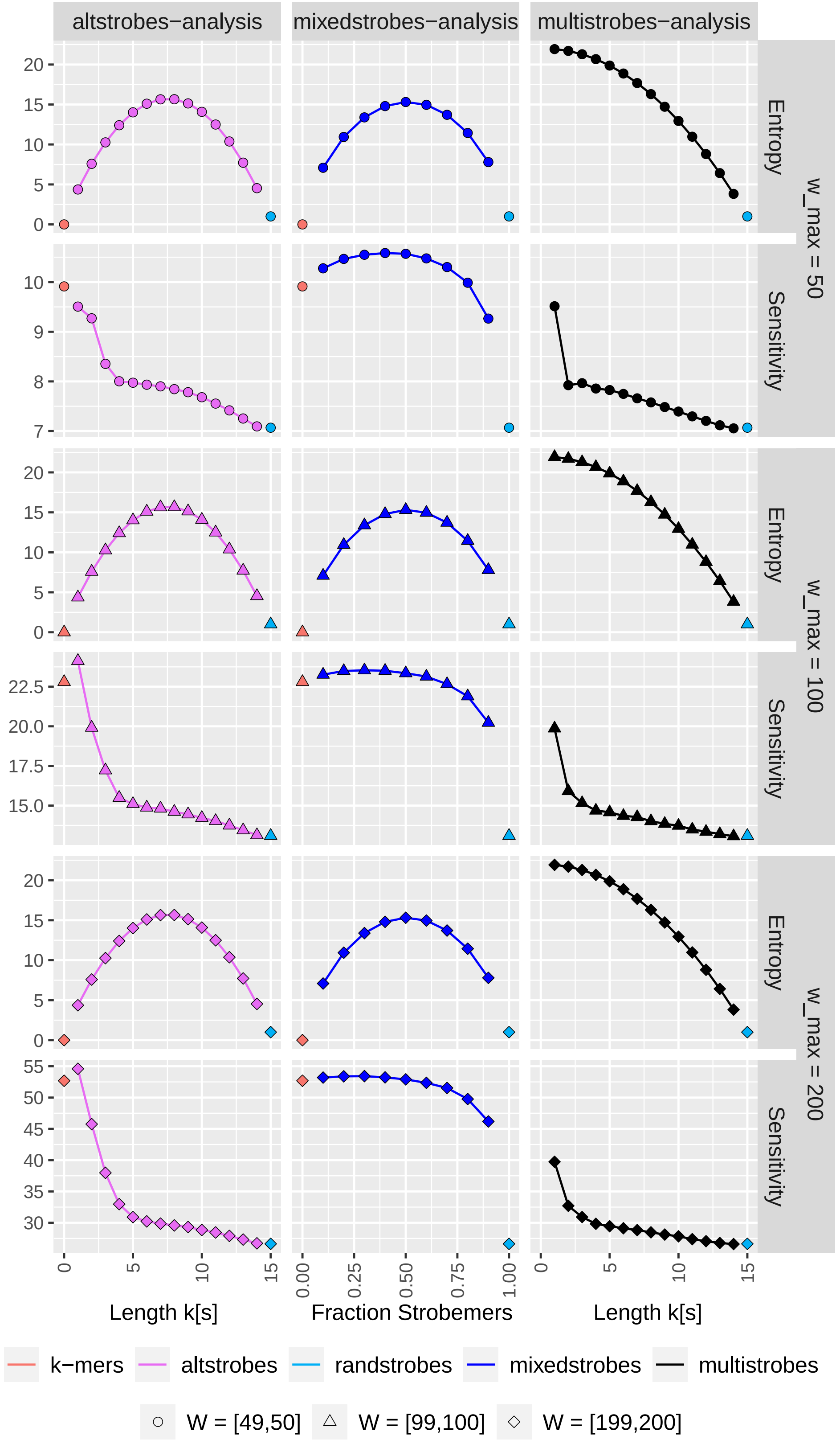


**Figure S2. Simulations showing entropies and sensitivity estimates for mixedstrobes, altstrobes, and multistrobes with very narrow window sizes ($w_{min}$,$w_{max}$) of (49,50), (99,100), and (199,200)**
Panel Entropy shows $H(\mathbf{X})$ for $k$-mers, randstrobes, altstrobes (for different $(k_s, k_l)$), mixedstrobes (for different $q$) and multistrobes (for different $(k_s, k_l)$). Panel Sensitivity shows $P(N_m(30, 64) > 0)$ summed over $m \in [1, 60]$ for various small window sizes.

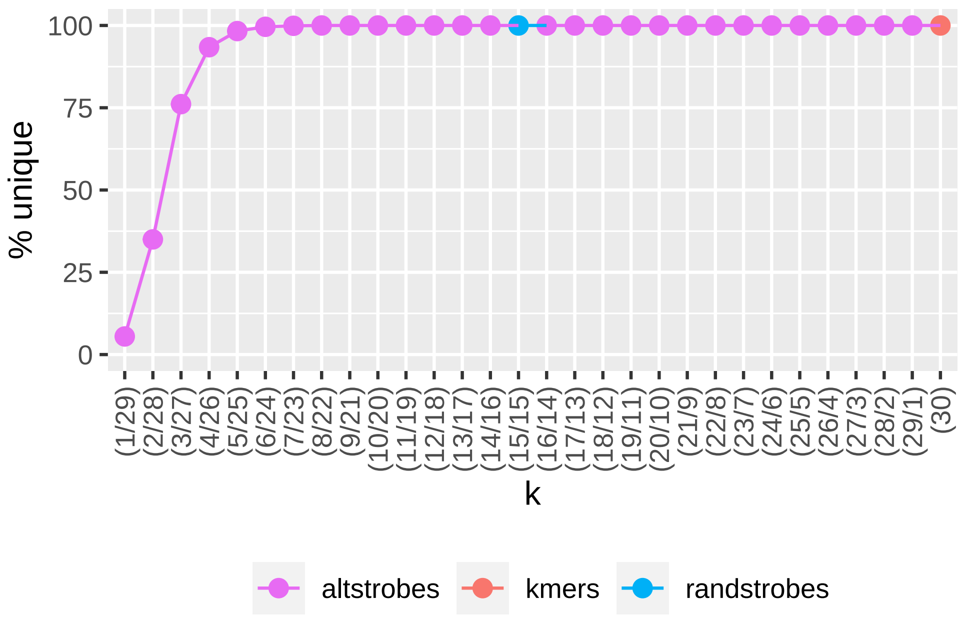


**Figure S3. Fraction of unique seeds for different altstrobe combinations in a region of 10,000nt.**
For a given altstrobe combination $(k_1/k_2)$ in the plot, the uniqueness of seeds was computed with parameters $(2, k_1/k_2, 25, 50)$ over simulated sequences of length 10.000nt. Note that all altstrobe seeds were sampled as $k_1/k_2$, which means that altstrobes(1/29) seeds only contain (1/29) seeds and no (29/1) seeds as opposed to all other altstrobe analysis in this manuscript.

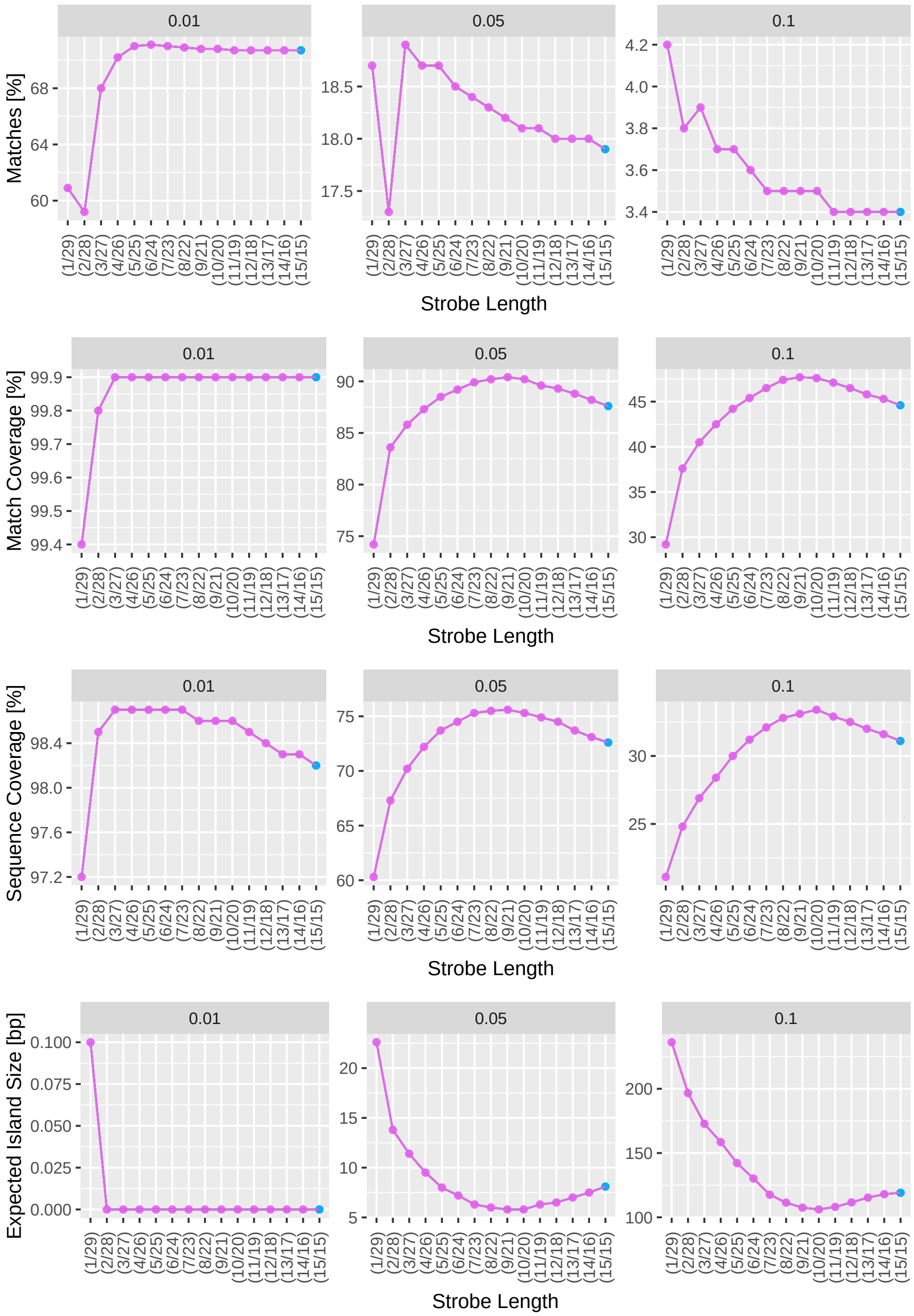


**Figure S4. Match Statistics for altstrobes with different strobe length combinations.**
For a given strobe length combination and mutation rate in the plot, 1,000 random sequences of length 10,000nt with randomly updated hash tables were generated. To guarantee an equal number of long-short and short-long altstrobe combinations even for very short strobes (see concerns prompted in section 1. Construction of Altstrobes), altstrobes were constructed with a modified altstrobe implementation. To this end, reference sequences were seeded with both altstrobes possibilities for each position and the decision about whether a short-long or a long-short query seed should be constructed was based on the longer strobe ensuring a large enough hash space to neglect systematic bias.

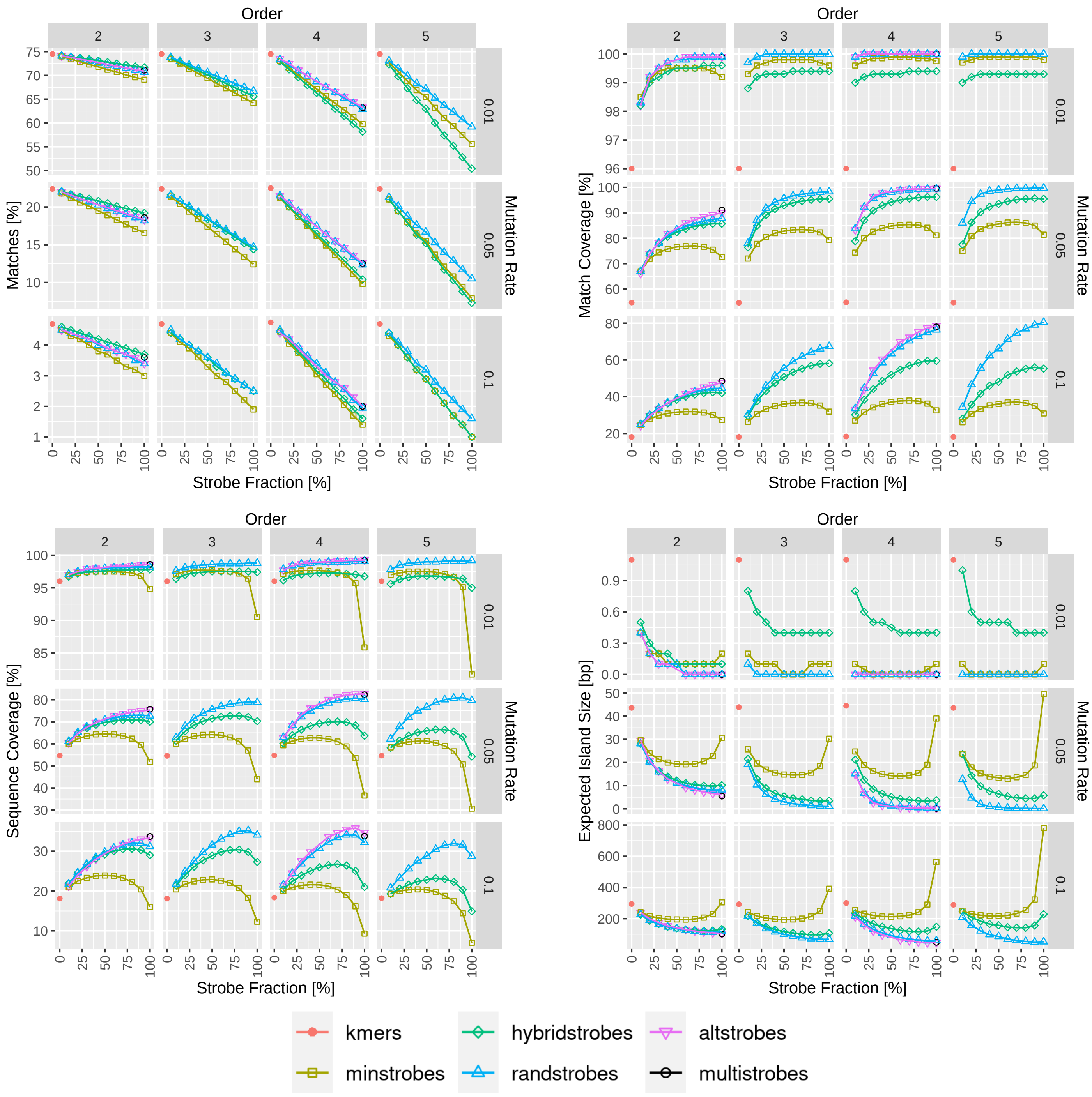


**Figure S5. Match Statistics for $k$-mers, strobemers, altstrobes, multistrobes and mixedstrobes of orders 2 - 5.**
1,000 random DNA sequences of 10,000 nucleotides were generated and subsequently mutated to obtain references and queries. Using these sequences, seeds were sampled with strobemer fractions from 0% ($k$-mers) to 100% (pure strobemers), downstream windows set to [25,50] and all strobes combined adding up to equal length subsequences of size 30 for better comparison. Hence, the strobemer settings were as follows: (2,15,25,50,$q$), (3,10,25,50,$q$), and (5,6,25,50,$q$), whereby mixedstrobes were sampled with a strobe fraction $q$ ranging from 0% ($k$-mers) to 100% (strobemers) and a step size of 10% ($f(0, 100, 10) = \{10k \mid k \in \{0, 10, ..., 100\}\}$). For strobemers of order 4, seeds of lengths 28 and 32 were seeded and the average (mean) plotted for each metrics to ensure similar sizes of subsequences between the protocols. Altstrobes were seeded with strobes of length $k_s = 10/k_l = 20$ (2,10,20,25,50,$q$), and $k_s = 5/k_l = 10$ (4,5,10,25,50,$q$), respectively. Multistrobes were seeded as (2,5,25,25,50) and (4,5,10,25,50).

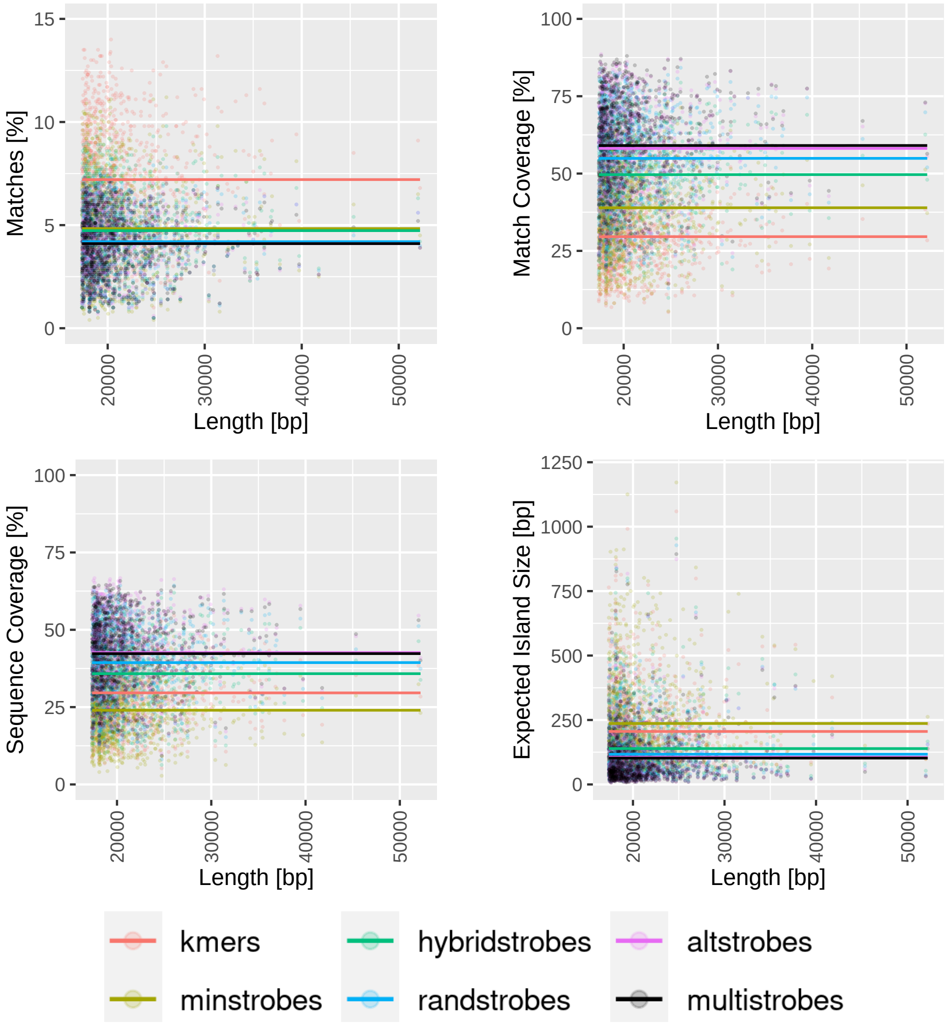


**Figure S6. Comparison between $k$-mers, strobemers, altstrobes and multistrobes when mapping genomic ONT reads for reads of different lengths (x-axis).**

The *E.coli* reads were split up in long disjoint segments of 2,000nt. Next, the segments were seeded with $k$-mers, strobemers and altstrobes, downstream windows set to [25,50] and all strobes combined adding up to equal length subsequences of size 30 for better comparison. Then for each segment, the collinear solution of raw hits was computed to subsequently quantify number of matches, match coverage, sequence coverage and expected island size for each read. Each dot represents one read while the line displays a smoothed conditional mean (GAM curve with cubic spline: $y \sim s(x, bs = "cs")$).

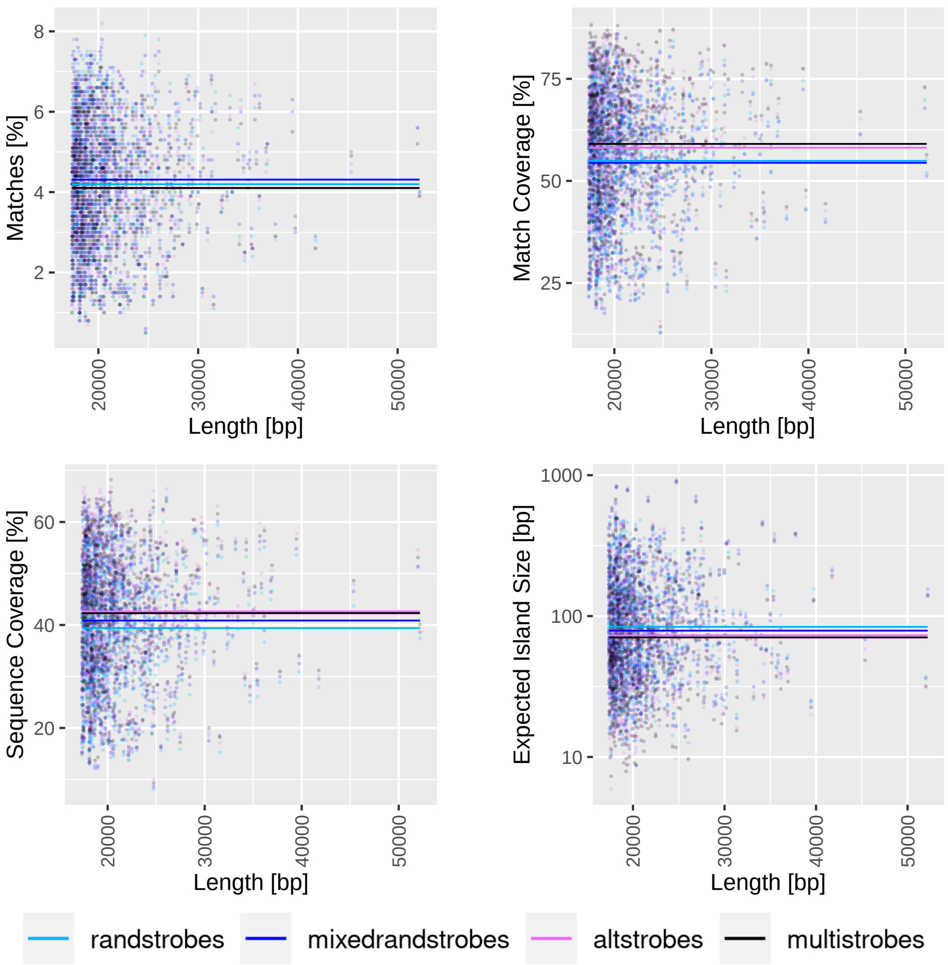


**Figure S7. Comparison between randstrobes, mixedrandstrobes, altstrobes and multistrobes when mapping genomic ONT reads for reads of different lengths (x-axis).**

The *E.coli* reads were split up in long disjoint segments of 2,000nt. Next, the segments were seeded with randstrobes (2,15,25,50), mixedrandstrobes (2,15,25,50,0.8), altstrobes (2,10,20,25,50) and multistrobes (2,5,25,25,50). For each segment, the collinear solution of raw hits was computed to subsequently quantify number of matches, match coverage, sequence coverage and expected island size for each read. For each read, the matching metrics from mixedrandstrobes (blue), altstrobes (pink) and multistrobes (black) were subsequently normalized by randstrobes (turkis) for better visualisation. Multistrobes and altstrobes perform best indicated by similar number of matches, higher sequence and match coverage as well as lower gap size. Mixedrandstrobes perform better than randstrobes for all metrics besides match coverage where mixedrandstrobes perform roughly 5% worse. Each dot represents one read while the line displays a smoothed conditional mean (GAM curve with cubic spline: $y \sim s(x, bs = "cs"))$.

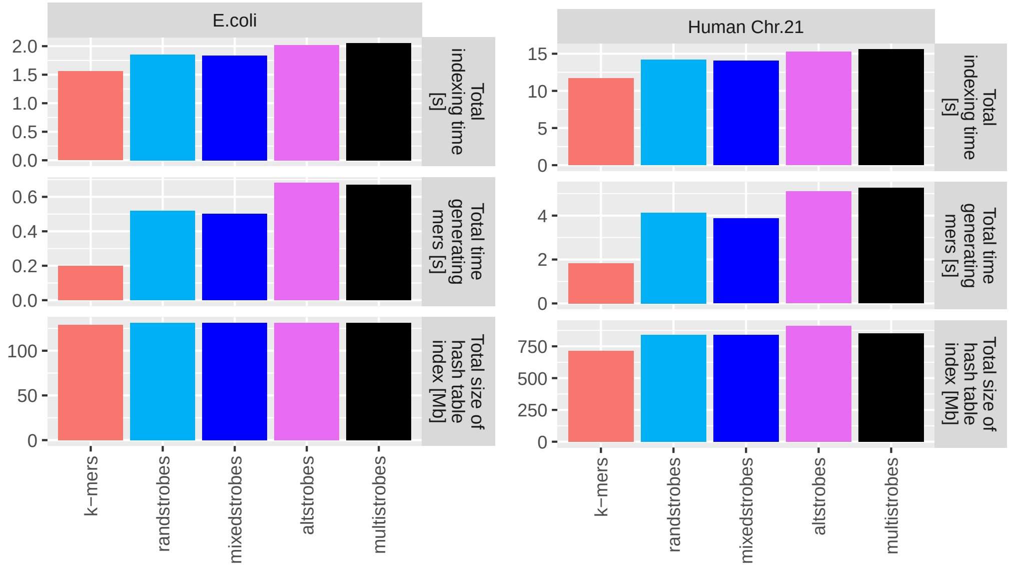


**(a)** *E.coli* **(b)** *Homo Sapiens* Chr.21

**Figure S8. StrobeMap Benchmarking**
To benchmark our two new seeding techniques, chromosome 21 of the GRCh38 human genome assembly and one *E.coli* genome (assembly GCA 003018575.1 ASM301857v1) were indexed with $k$-mers ($k = 30$), randstrobes $(2, 15, 25, 50)$, mixedstrobes $(2, 15, 25, 50, 0.8)$, altstrobes $(2, 10, 20, 25, 50)$ and multistrobes $(2, 5, 25, 25, 50)$ using StrobeMap. All experiments were repeated 10 times and the average (mean) was computed to account for variance in computer processing speed.

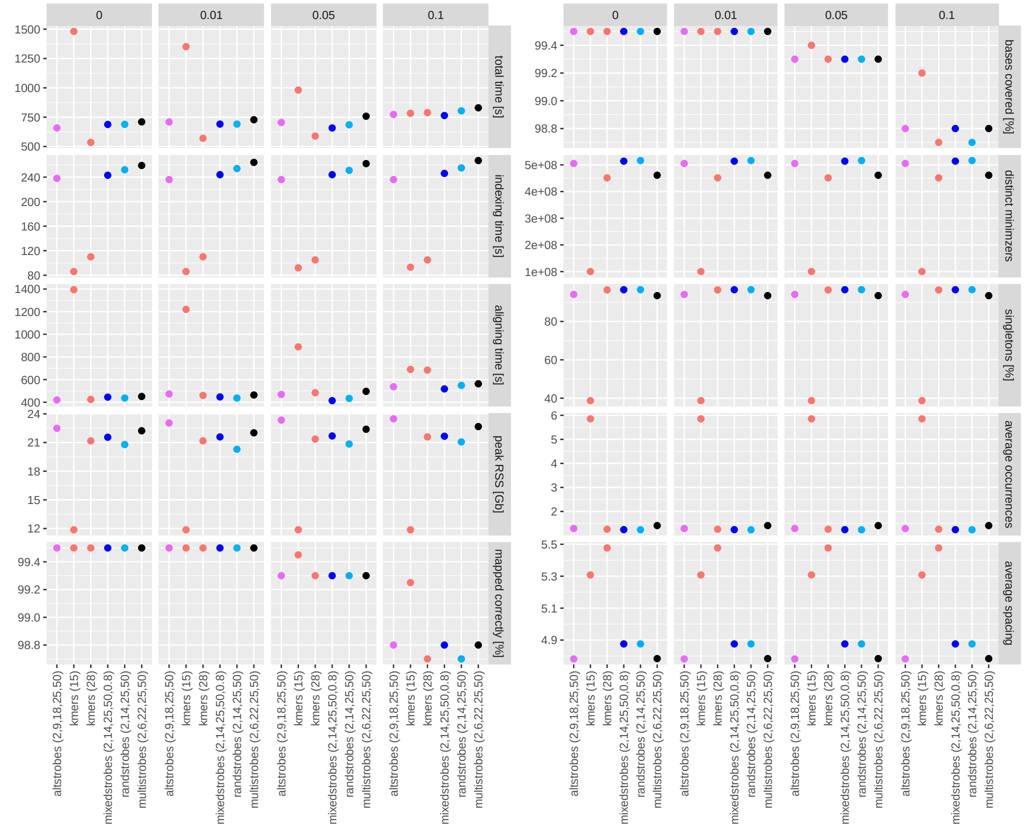


**Figure S9. Minimap2 Benchmarking.**
100.000 sequences of lengths 10.000nt without any incompletely specified bases (e.g. ”n”) were selected from random positions of the CHM13v2.0 human assembly and mutated (insertions, deletions and base substitutions) with mutation frequencies of 0%, 1%, 5% and 10%. The selected reads were mapped and aligned back to the reference using $k$-mers ($k$ = 15 and 28), altstrobes $(2, 9, 18, 25, 50)$, randstrobes $(2, 14, 25, 50)$, multistrobes $(2, 5, 23, 25, 50)$ and mixedstrobes $(2, 14, 25, 40, 0.8)$. All experiments were repeated 5 times and the average (mean) taken to account for variance in computer processing speed.

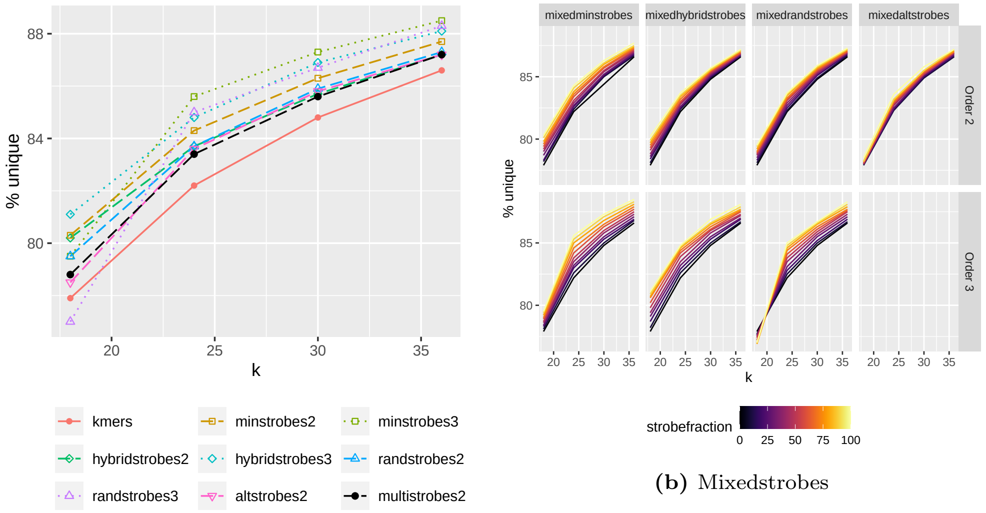


**(a)** Overview

**(b)** Mixedstrobes

**Figure S10. Fraction of unique seeds for different strobemer seeding techniques and fractions on Human chromosome 21.**
Chromosome 21 of the human GRCh38 assembly was seeded with $k$-mers, strobemers of orders 2 (2,$k$/2,25,50,$q$) and 3 (3,$k$/3,25,50,$q$) as well as altstrobes of order 2 (2,($k$/3),(2$k$/3),25,50,$q$). Mulitstrobes was seeded with (2,5,$k$-5,25,50). In panel B, the numbers of extracted nucleotides ($k$=30) is the same for all seeding techniques for fair comparison.

w = 1

| Mutation Rate | | 0.01 | | | | 0.05 | | | | 0.1 | | | |
|---|---|---|---|---|---|---|---|---|---|---|---|---|---|
| Seeding | Settings | m | mc | sc | e | m | mc | sc | e | m | mc | sc | e |
| $k$-mers | 30 | **74.5** | 96.0 | 96.0 | 1.1 | **22.4** | 54.7 | 54.7 | 43.6 | **4.7** | 18.1 | 18.1 | 293.8 |
| spaced $k$-mers | dense | 67.6 | 95.6 | 96.2 | 1.5 | 13.8 | 53.9 | 50.9 | 65.7 | 1.8 | 16.3 | 14.2 | 481.0 |
| spaced $k$-mers | sparse | 50.5 | 89.8 | 87.8 | 11.1 | 3.5 | 26.8 | 21.4 | 493.5 | 0.1 | 3.6 | 2.1 | 4120.8 |
| altstrobes | (2,10,20,25,50) | 70.9 | **99.9** | **98.6** | **0.0** | 18.3 | 90.0 | 75.3 | 6.1 | 3.4 | 47.7 | 33.3 | 104.4 |
| randstrobes | (2,15,25,50) | 70.7 | **99.9** | 98.2 | **0.0** | 18.2 | 87.8 | 72.7 | 8.2 | 3.4 | 44.6 | 21.2 | 118.1 |
| hybridstrobes | (2,15,25,50) | 71.7 | 99.6 | 97.8 | 0.1 | 19.2 | 85.7 | 70.0 | 10.2 | 3.7 | 42.0 | 29.0 | 132.7 |
| minstrobes | (2,15,25,50) | 69.1 | 99.2 | 94.8 | 0.2 | 16.6 | 72.6 | 51.9 | 30.7 | 3.0 | 27.4 | 16.0 | 304.0 |
| mixedstrobes | (2,15,25,50,0.8) | 71.5 | **99.9** | 98.2 | 0.1 | 19.0 | 86.8 | 72.9 | 8.4 | 3.7 | 43.8 | 32.1 | 113.4 |
| multistrobes | (2,5,25,25,50) | 71.1 | **99.9** | **98.6** | **0.0** | 18.6 | **91.1** | **75.7** | **5.5** | 3.6 | **48.5** | **33.7** | **101.2** |

w = 10

| Mutation Rate | | 0.01 | | | | 0.05 | | | | 0.1 | | | |
|---|---|---|---|---|---|---|---|---|---|---|---|---|---|
| Seeding | Settings | m | mc | sc | e | m | mc | sc | e | m | mc | sc | e |
| $k$-mers | 30 | **73.2** | 90.3 | 90.3 | 2.6 | **20.7** | 42.8 | 42.8 | 73.2 | **3.9** | 11.4 | 11.4 | 501.8 |
| spaced $k$-mers | dense | 65.5 | 90.9 | 87.3 | 3.9 | 12.1 | 36.5 | 30.9 | 147.8 | 1.4 | 6.9 | 5.3 | 1265.4 |
| spaced $k$-mers | sparse | 47.9 | 84.9 | 74.4 | 17.0 | 2.7 | 16.7 | 9.5 | 945.9 | 0.1 | 1.2 | 0.5 | 7140.2 |
| altstrobes | (2,10,20,25,50) | 69.7 | 98.5 | 92.3 | **0.4** | 17.1 | 62.7 | 46.0 | 46.9 | 3.0 | 19.2 | 12.1 | 423.3 |
| randstrobes | (2,15,25,50) | 69.6 | 98.4 | 92.3 | 0.5 | 16.8 | 62.3 | 45.9 | 48.7 | 2.9 | 18.5 | 11.8 | 451.6 |
| hybridstrobes | (2,15,25,50) | 68.4 | 97.3 | 90.2 | 1.2 | 15.8 | 58.6 | 42.5 | 58.0 | 2.6 | 16.8 | 10.5 | 506.6 |
| minstrobes | (2,15,25,50) | 68.0 | 98.1 | 87.8 | 0.7 | 15.2 | 58.8 | 37.1 | 67.4 | 2.5 | 16.5 | 8.8 | 611.2 |
| mixedstrobes | (2,15,25,50,0.8) | 70.5 | 98.0 | 92.2 | 0.7 | 17.8 | 60.4 | 46.8 | 49.2 | 3.2 | 17.8 | 12.4 | 425.6 |
| multistrobes | (2,5,25,25,50) | 70.3 | **98.6** | **92.4** | **0.4** | 17.2 | **64.2** | **47.0** | **44.1** | 3.3 | **20.2** | **12.7** | **404.2** |

w = 20

| Mutation Rate | | 0.01 | | | | 0.05 | | | | 0.1 | | | |
|---|---|---|---|---|---|---|---|---|---|---|---|---|---|
| Seeding | Settings | m | mc | sc | e | m | mc | sc | e | m | mc | sc | e |
| $k$-mers | 30 | **71.8** | 84.2 | 84.2 | 5.8 | **19.3** | 33.2 | 33.2 | 111.1 | **3.6** | 7.8 | 7.8 | 737.4 |
| spaced $k$-mers | dense | 64.4 | 85.0 | 77.9 | 8.0 | 11.4 | 27.9 | 21.9 | 221.3 | 1.4 | 4.5 | 3.3 | 1931.3 |
| spaced $k$-mers | sparse | 47.3 | 79.9 | 62.2 | 26.6 | 2.6 | 12.2 | 5.9 | 1256.4 | 0.1 | 0.7 | 0.3 | 8242.6 |
| altstrobes | (2,10,20,25,50) | 68.9 | 94.8 | 83.6 | 2.3 | 16.4 | 47.0 | 32.1 | 98.5 | 2.9 | 11.6 | 7.1 | 766.8 |
| randstrobes | (2,15,25,50) | 68.3 | 94.8 | **83.9** | 2.4 | 15.6 | 46.0 | 31.2 | 101.3 | 2.7 | 11.0 | 6.7 | 804.9 |
| hybridstrobes | (2,15,25,50) | 66.2 | 92.6 | 81.8 | 3.8 | 13.8 | 41.1 | 28.1 | 124.8 | 2.2 | 9.2 | 5.6 | 993.7 |
| minstrobes | (2,15,25,50) | 66.7 | 95.3 | 75.8 | 2.6 | 14.0 | 45.4 | 25.6 | 129.3 | 2.2 | 10.4 | 5.2 | 1020.3 |
| mixedstrobes | (2,15,25,50,0.8) | 69.3 | 93.3 | 83.8 | 2.9 | 16.6 | 44.3 | 32.4 | 99.0 | 2.9 | 10.6 | 7.2 | 768.5 |
| multistrobes | (2,5,25,25,50) | 69.8 | **95.0** | 83.6 | **2.2** | 17.2 | **48.3** | **32.8** | **93.9** | 3.1 | **12.2** | **7.4** | **722.8** |

**Table S1: Match statistics for simulated sequences ($L = 10000$)) under different sampling protocols under mutations rates of 0.01, 0.05, 0.1 using minimizer thinning with $w = 1$ (no thinning), $w = 10$, and $w = 20$.**
Here, m denotes the number of matches as a percentage of the total number of extracted subsequences for the protocol, sc (sequence coverage) and mc (match coverage) is shown as the percentage of the total sequence length, and E is the expected island size. Boldfaced values indicate the most desirable result across protocols for each of the match statistics.
